# Cross-tissue isoform switches by human transcription factors suggest a widespread regulatory mechanism

**DOI:** 10.1101/2024.11.28.625692

**Authors:** Sviatoslav Sidorov, Koustav Pal, Boris Lenhard, Nicholas M. Luscombe

## Abstract

Many human genes, including transcription factor (TF) genes, produce alternative coding isoforms. Although some alternative coding TF isoforms have been experimentally studied, a genome-wide characterisation of all such isoforms in normal human physiology is still lacking. Here, we integrate RNA sequencing and protein domain data to reveal the expression levels, tissue specificity and putative functional classes of alternative coding isoforms of human TF genes. We found that isoforms lacking a DNA-binding domain (DBD) are annotated for almost a third of known TF genes, comprise 17% of all annotated TF isoforms and likely reverse the regulatory effects of DBD-containing isoforms. Considering all coding TF isoforms, we uncovered 536 switches between the highest-expressed alternative isoforms produced by 360 TF genes across 43 adult tissues. We predicted functional consequences of these switches using a functional annotation of structural domains present in switching isoforms. Furthermore, we found 45 isoform switches between the brain and non-brain tissues and interpreted six of these switches in detail. Finally, we assembled a literature-based compendium of human TF genes with experimentally studied alternative isoforms. In total, our results suggest that the switches of alternative coding TF isoforms between adult tissues represent a widespread, yet understudied, functional mechanism.

## Introduction

A gene may produce more than one coding isoform using alternative splicing, as well as alternative transcription start and termination sites. About 95% of human genes undergo alternative splicing (1, 2), which potentiates large-scale proteome diversification. Although the extent of translation of alternative mRNA isoforms has been actively debated (3–15), many studies have shown that alternative splicing produces proteins with different functions and regulates molecular processes. For instance, alternative splicing affects protein-protein (16, 17), protein-DNA (18, 19) and other protein interactions (reviewed in (20)), as well as cellular signalling (21).

Alternative transcription initiation and termination further increase isoform diversity. Shabalina and colleagues demonstrated that the contribution of these mechanisms to the diversity of mammalian mRNA and protein isoforms is even greater than the one of alternative splicing (22). In addition, Reyes and Huber have shown that alternative transcription initiation and termination generate the most part of tissue-specific exon usage, although may not lead to considerable protein diversification (23). However, production of alternative mRNA isoforms may still regulate the level of translation (24) and, in general, has a broad range of consequences beyond increasing proteome complexity (reviewed in (25)).

Alternative isoform production by transcription factor genes (TF genes) is particularly intriguing, for it may influence transcription regulation genome-wide, during development or between different tissues, in health and disease. Alternative coding isoforms of specific TF genes have been detected and examined in various organisms, including plants (reviewed in (26, 27)), fungi (28–35), worms (36–39), insects (40–46) and mammals (47–54) and were shown to regulate key physiological processes. The roles of coding alternative isoforms of TF genes (hereafter, TF isoforms) have also been studied in development ((55); early results reviewed in (56)) and cancer (reviewed in (57)).

While experimental investigations of single TF isoforms are key to our understanding of TF isoform biology, such studies are laborious and still far from uncovering functions of alternative isoforms of all TF genes in a whole eukaryotic genome. On the other hand, recent groundbreaking efforts to build genome-wide atlases of human TF isoforms (58, 59) were not, by design, aimed at a functional characterization of every single isoform. As a compromise between the two approaches, a mid-scale experimental study of human TF isoforms by Lambourne and colleagues, described in their recent preprint (60), tested the DNA-binding, protein-binding and transcription regulation activities, as well as the subcellular localization and condensate formation, of a total of 693 isoforms produced by 246 TF genes in the brain, liver and heart. Although a major step forward, this study assayed only about 17% (693/4,144) of all coding TF isoforms (according to Lambourne and co-author’s estimation of the total number of such isoforms) and did not cover other possible functions of TF isoforms, such as ligand binding (60).

In contrast to experimental approaches, computational analyses enable genome-wide isoform studies. Early computational publications revealed that some TF domains are preferentially removed by alternative splicing and that a larger fraction of TF genes, in comparison to non-TF genes, undergo alternative splicing in the human and mouse (61–64). More recently, Soto and colleagues created TFRegDB, a publicly available service that allows scanning transcript sequences for >900 effector domains experimentally verified in ∼600 human TFs (65). However, in these studies, the authors did not predict functions of human TF isoforms. In contrast, a plethora of computational methods that predict isoform functions genome-wide summarise these functions only at the level of Gene Ontology terms and either do not take protein sequence and functional domains into account (66–73), or cannot incorporate current knowledge about protein domain functions (74–78). Therefore, a genome-wide study of human TF isoforms in normal physiology, based on the current knowledge about functional TF domains, is still lacking.

Here, we present a genome-wide computational survey of protein-coding alternative isoforms of human TF genes. We found that isoforms lacking a DNA-binding domain (DBD^−^ isoforms) comprise 17% of all TF isoforms and are annotated for almost a third of all TF genes. Moreover, 69% of DBD^−^ isoforms are expressed in at least one adult tissue. Intriguingly, we uncovered a subset of DBD^−^ isoforms present in a broad range of tissues and highly expressed in at least some of them, which suggests functional relevance. Furthermore, based on a functional catalogue of non-DNA-binding domains and a catalogue of previously characterised TF isoforms that we compiled for the present study, we predicted functional classes of DBD^−^ isoforms and inferred that they mainly reverse the regulation effects of DBD-containing (DBD^+^) isoforms. Moreover, considering all expressed TF isoforms, we revealed 536 switches between the highest-expressed (major) alternative isoforms produced by 360 TF genes across 43 adult human tissues and predicted the general functional consequences of these switches based on the functional annotation of hundreds of domain types. Our unsupervised clustering of the obtained switches uncovered 45 switches between the brain and non-brain tissues, and we interpreted six of these switches in detail. In summary, our computational analyses dissect, for the first time, a genome-wide set of alternative protein-coding isoforms of human TF genes and suggest that switches between these isoforms comprise an unexplored layer of transcription regulation.

## Material and Methods

### Selection of protein isoforms of human TF genes

We obtained a list of all human TF genes from the HumanTFs database (79), version 1.01, and downloaded sequences of their protein isoforms from the Ensembl database (80) release 99. We retained isoforms with Transcript Support Level (TSL) tags TSL1 and TSL2, as well as isoforms with TSL = NA and isoforms with any TSL but with the Matched Annotation from NCBI and EMBL-EBI (MANE) Select tag. We retained isoforms with TSL = NA because they did not undergo TSL analysis in Ensembl due to being produced by leukocyte antigens, immunoglobulin genes, T-cell receptor genes or pseudogenes, or due to containing a single exon (https://www.ensembl.org/info/genome/genebuild/transcript_quality_tags.html#tsl) and therefore are not necessarily of a low quality. Isoforms with the MANE Select tag are identical in the Ensembl and NCBI RefSeq (81) human genome annotations and are independently chosen by these consortia as the most biologically relevant for a gene (https://www.ensembl.org/info/genome/genebuild/transcript_quality_tags.html#mane). We therefore considered isoforms with the MANE Select tag as correct, independently of their TSL.

### Domain annotation in protein isoforms

We used InterProScan 5 (82), version 5.39-77.0, to search for domains in the protein sequences of all TF gene isoforms initially obtained from the Ensembl (we selected isoforms by annotation quality later on; see *Definition of TF gene families*). For each InterPro entry match, we calculated its **integrated coordinates** as the minimal and maximal signature coordinates across all signatures matched within the entry. In the calculation, we did not consider matched signatures from non-domain databases PANTHER and PIRSF. Next, we deduplicated overlapping entry matches by classifying all entry matches into "general" and "specific" and then removing the "general matches". We called a match of an InterPro entry *E* in an isoform *P* **general**, if *E* is a Domain or a Family and at least one of its descendant entries also matched *P* and overlapped with the match of *E* by at least one amino acid. Otherwise, we called a match of an InterPro entry **specific**. We classified InterPro entry matches into general and specific with custom Bash and Python scripts and R packages *yaml* (Garbett et al., 2022), version 2.3.5, and *data.tree* (Glur, 2020), version 1.0.0.

Because entry matches may overlap, we deduplicated them by ranking entry matches according to their type in the following order: Domain; Family matches with domain signatures from Pfam, SMART, PrositeProfiles or CDD (we defined a separate Domain-Family type for such entry matches); Homologous Superfamily; Site; Repeat; Family (Family matches without domain signatures). This ranking is based on the fact that the Domain and Family-Domain entries, by definition, represent domains; Homologous Superfamily entries also consist of signatures that represent domains: CATH-Gene3D and/or SUPERFAMILY; Site and Repeat matches represent short functional peptides within a protein (https://www.ebi.ac.uk/training/online/courses/interpro-functional-and-structural-analysis/wha <u>t-is-an-interpro-entry/interpro-entry-types/</u>). Next we excluded matches of a lower rank that overlapped with specific matches of a higher rank by integrated coordinates. Also, we manually deduplicated overlapping specific matches of each type, apart from Domain that we processed separately later. If a general match did not overlap with a specific match after these steps, we redefined the general match as specific.

After deduplicating matched InterPro entries, we selected one matched signature in each entry (specific or general) and removed other signatures. Specifically, we prioritised signatures in the following way: Pfam, SMART, ProSiteProfiles, CDD, SUPERFAMILY, Gene3D, PRINTS, ProSitePatterns, PIRSF, PANTHER, starting from domain signatures (Pfam to Gene3D), followed by conserved motif signatures (PRINTS and ProSitePatterns) and protein family signatures (PIRSF and PANTHER). Using this ranking, we retained in each matched entry only the signature with the highest rank. Additionally, for some entry matches we manually corrected the choice of a representative signature. For example, for C2H2 ZF and CCCH ZF domain entry matches, we chose SMART signatures, if both SMART and Pfam signatures matched, because, based on a visual inspection, SMART signatures represent these zinc finger domains more adequately. Also, if a domain was represented by sequential matches of Gene3D, PRINTS or PROSITE pattern signatures and was not a C2H2 ZF array, we merged these signature matches to obtain a continuous domain representation.

### Classification of domains annotated in protein isoforms

Using a custom Python script, we initially classified all InterPro entry matches (specific and general) as "DNA-related" (domains with at least one mention of the word "DNA" in their InterPro entry name, description or associated Gene Ontology terms) and "Non-DBD" (other domains). We removed Family matches from this preliminary classification, because these matches do not represent domains. Next, within the DNA-related category, we manually separated "DBD", "Other", "Unknown" and "Unclear" categories according to the following definitions:

1. The **DBD category** includes DNA-binding domains, described as such in the InterPro.
2. The **Other category** includes domains with functions that are different from DNA binding, according to the InterPro.
3. The **Unknown category** includes domains whose functions are as yet unknown, according to the InterPro.
4. The **Unclear category** includes domains whose functions are not mentioned in their InterPro entry description.

In this way, we separated DBDs from other domains and formed a list of non-DBDs (represented by the union of the Other and Non-DBD categories) for further classification.

### Definition of TF gene families

Before using the obtained DBD annotation of the protein isoforms to define TF gene families, we corrected some DBD entry matches **(Table S11)**. We also merged disrupted DBD matches in several isoforms. Additionally, we restored the following DBD signature matches lost in filtering: AT hook (IPR017956, SMART) in BAZ2A (isoforms ENST00000379441, ENST00000549884, ENST00000551812); C2H2C ZF (IPR002515, Pfam) in ST18 (isoform ENST00000276480) and C2H2C ZF (IPR002515, Pfam) in MYT1L (isoform ENSG00000186487). Next, we removed general DBD matches, either by redefining them as specific (if they did not overlap with any other matches) or by deleting them from the annotation (if they overlapped specific matches). Furthermore, we removed redundant (specific) DBD matches by retaining the longest match in each set of overlapping DBD matches (or by manually selecting a match of a more specifically named/described InterPro entry in the case of matches of equal length). After these refinement steps, we grouped matched InterPro entries corresponding to the same DBD type and defined **a TF gene family** as a group of TF genes that encode the same unique combination of DBD types. We removed genes with an uncertain TF status, according to the HumanTFs database and genes without a well-known DBD. For the excluded TF genes, we defined two technical categories: “Excluded” (TFs that we excluded from the further analysis during the curation described above) and “Unknown” (genes that do not have a known DBD, according to both the HumanTFs database and our analysis).

After obtaining the final assignment of TF genes to families, we filtered isoforms by the transcript support level and the presence of the MANE Select tag (see *Selection of protein isoforms of human TF genes*).

### Functional annotation of non-DNA-binding domains and DBD^−^ isoforms

We used InterPro entry descriptions and extensively searched literature to predict functions of 262 unique InterPro entries corresponding to general and specific matches of non-DNA-binding domains (non-DBDs). We included general matches to obtain better understanding of non-DBD functions **(Table S5)**. To deduplicate non-DBD InterPro entry matches, we found those non-DBD matches that overlapped by ≥70% of their length with at least one other non-DBD match and retained one or both overlapping matches according to the following rules:

1. If the two matches represent the same domain, then retain the longer match.
2. If the two matches represent different domains, or it is unclear if they represent the same domain, then:

a. If the two matches overlap reciprocally by ≥70% of their lengths, then retain a more biologically relevant match or a match with a more detailed description in the InterPro.
b. Otherwise, retain both matches, as they may represent a combination of domains of two different types.

After the deduplication of non-DBD matches, we grouped DBD^−^ isoforms into general functional classes based on the functions of matched non-DBD combinations **(Table S6)** and examples of functional DBD^−^ isoforms found in literature.

### Isoform expression analysis

To estimate isoform expression across various human tissues, we used the Analysis Freeze dataset from the Genotype-Tissue Expression (GTEx) database (83), version 8. From this dataset, we selected tissues with at least 100 samples and removed “Cells – Cultured fibroblasts” and “Cells – EBV-transformed lymphocytes”, as these are cell lines; we also removed “Whole Blood” because it represents a very heterogeneous group of cells. For each of the 43 obtained tissues, we selected 50 samples at random. We normalised raw isoform expression counts across the 2,150 selected samples with the *estimateSizeFactors* function from the DESeq2 R package (84). We defined the **expression level of an isoform in a tissue** as its median expression level across the 50 samples randomly chosen for that tissue. We defined an isoform as **expressed in a tissue** if its expression level in that tissue is at least 5 DESeq2-normalised counts. Accordingly, we defined an isoform as **expressed** if it is expressed in at least one tissue. We defined the final set of expressed isoforms as the intersection of the set of annotated TF isoforms obtained from the Ensembl v99 with the set of expressed isoforms. Additionally, we defined the **buffered expression level of an isoform** as the mean of the top 20% of its expression levels across tissues in which the isoform is expressed.

For the calculation of the tau score which quantifies the tissue specificity of isoform expression (85), we subtracted the minimal expression cutoff of 5 DESeq2-normalised counts from the expression level of each isoform in each tissue and replaced obtained negative values with zero. In this way, we retained a continuous scale of expression values. Additionally, by replacing the expression values under the minimal expression cutoff with zero, we avoided affecting tissue specificity scores of isoforms by their negligible expression levels in tissues where they are not counted as expressed. Because the tau score is very sensitive to high isoform expression in particular tissues, even if the isoform is expressed uniformly across the majority of other tissues, we log2-scaled the obtained counts for the score calculation to decrease the expression variation. We calculated the tau score according to its original definition (85):

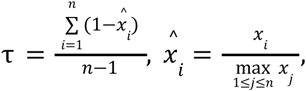

where 𝑥𝑖 is the expression level of an isoform in tissue 𝑖, and 𝑛 is the number of tissues.

To verify our use of the tau score as a measure of tissue specificity, we searched for DNA-binding subunits of general TFs in literature and intersected the list of the obtained subunits with the set of TFs that we used in our analysis. Additionally, we used single-cell gene-level transcription quantification data from the Human Protein Atlas (86) to find broadly expressed and tissue-specific TFs. Namely, for some of the tissues shown at https://www.proteinatlas.org/humanproteome/tissue, we selected the list of enriched genes and among those searched for TFs.

### Detection and analysis of major isoform switches

We defined **an expression level of an isoform in a tissue** (in transcripts per kilobase per million, or TPMs) as isoform’s median GTEx v8 TPM expression level across the 50 samples randomly selected for this tissue (see above). We defined **a major isoform of a gene** as an isoform with the highest TPM expression level among all isoforms of the gene in at least one tissue. In this definition, we used TPM expression levels because they allow comparing the expression levels of different isoforms within the same tissue. Of note, according to this definition, a gene may have more than one major isoform.

Based on the set of major isoforms, we detected major isoform switches. We defined **a switch between major isoforms *K* and *J* of the same TF gene** as an exchange of the isoform *K* expressed as major in one set of tissues for the isoform *J* expressed as major in another set of tissues, such that the difference between the expression levels of the two isoforms in any tissue from the two sets is at least 20% **(Figure 4A)**.

We summarised switches by assigning each TF gene that switches major isoforms to a category in each tissue, according to the following rules. Let a TF gene G produce two major isoforms, A and B, that switch between a set of tissues T_A_, where A dominates B, and a set of tissues T_B_, where B dominates A. Let *N*_A_ and *N*_B_ be the sizes of T_A_ and T_B_, respectively, such that *N*_A_ ≥ *N*_B_. Assign a category **a** (**b**) to G in tissues from T_A_ if A is DBD^+^ (DBD^−^). Likewise, assign a category **c** (**d**) to G in tissues from T_B_ if B is DBD^+^ (DBD^−^). For instance, if A is DBD^+^, then G will be assigned the category **a** in tissues from T_A_, and if B is DBD^−^, then G will be assigned the category **d** in tissues from T_B_. In all tissues not from T_A_∪T_B_, assign G a category **e**. Therefore, G is assigned the category **e** in all tissues where G is not expressed or where it is expressed but does not switch its major isoforms.

We organised the TF gene-category assignments into a matrix in which rows correspond to switches, columns correspond to tissues, and each cell contains a category assigned to a given TF gene that switches a given pair of its major isoforms. We used the ComplexHeatmap R package v2.14.0 (87) to cluster rows and columns of the heatmap by the Hamming distance between category vectors and to visualise the clustered heatmap.

We used the implementation of the hypergeometric test from the hypeR R package v1.14.0 (88) to test gene set overrepresentations. We adjusted raw p-values for multiple testing using the Benjamini-Hochberg procedure implemented in the *p.adjust* base R function.

### Code availability

Source code for the described data processing and analysis can be found at https://github.com/ComputationalRegulatoryGenomicsICL/tf-isoforms-publication.

## Results

### Known examples of TF isoforms provide the basis for a genome-wide analysis

To build an interpretation basis for our TF isoform analysis, we studied literature and compiled a collection of TF genes that produce characterised alternative protein isoforms with functional differences explained through differences in structural domains **(Table S1)**. These examples include the differential presence of DNA-binding domains (DBDs), transcription regulation domains and other domains **(Figure 1, S1)**. Thus, the differential inclusion of a particular DBD (out of several DBDs), as well as a full or partial exchange of one DBD sequence for another, may lead to a change in the DNA binding affinity **(Figure 1A)**. The differential presence of a single DBD may result in opposite transcription regulation effects through the sequestration of a co-binding TF or cofactors with a dimerization or transcription regulation (cofactor binding) domain **(Figure 1B-C)**. The regulation effect can also be reversed by the differential presence of a transcription activation **(Figure 1D)** or repression **(Figure 1E)** domain, while differences in the sequence of the transcription regulation domain may lead to changes in the specificity **(Figure 1F)** or strength **(Figure 1G)** of the regulation. Finally, functional changes in domains of other types **(Figure S1A-D)**, as well as more complex, but functionally important, differences in transcription regulation domains **(Figure S1E-G)**, are also known. Overall, we use these experimentally studied examples to predict functional classes of the isoforms of human TF genes. Importantly, the fact that dozens of human TF genes produce functional alternative isoforms signifies the necessity of studying human TF isoforms genome-wide.

**Figure 1.**
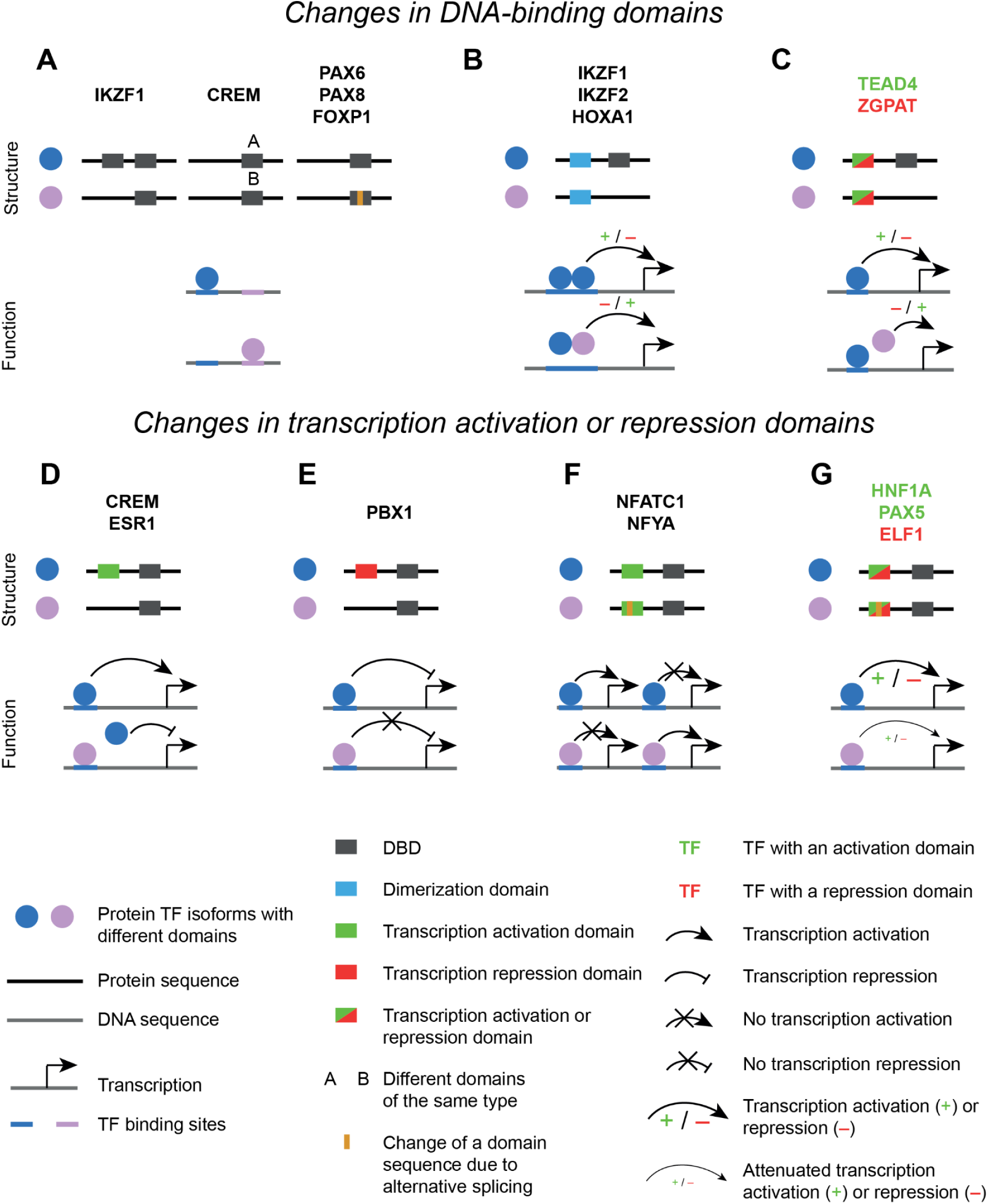
Differences in domains between alternative isoforms modulate the function of a TF gene. **(A-C**) Differences in DBDs. **(D-G)** Differences in transcription regulation domains. See **Table S1** for details.

In addition to TF genes listed in **Figure 1 and S1 (Table S1)**, we present a collection of another 70 TF genes (or groups of closely related TF genes) whose alternative isoforms were studied to various extent **(Table S2)**. We anticipate that our total collection of 92 TF genes and groups of TF genes will help the future studies of the human TF isoforms.

### Human TF genes widely produce DBD-lacking isoforms

TFs are proteins that bind DNA and regulate transcription. By this definition, a protein-coding isoform produced by a TF gene but missing a DBD cannot act as a TF, presenting the most dramatic example of a domain change effect upon the function of a TF gene. Therefore, we started our analysis of human TF isoforms by studying the differential inclusion of a DBD. To this end, we used InterProScan (82) to delineate known structural domains in 4,134 protein sequences of coding TF isoforms with a high-quality annotation in Ensembl (80) **(see Table S3 as a Supplementary Data File)**. We found that 17% (705) of the isoforms lack a DBD, and we denoted such isoforms DBD^−^. These isoforms were annotated for 27.5% (415/1,512) of high-confidence human TF genes **(Figure 2A)** that we selected for this study from the HumanTFs database (see **Material and Methods** for details) (79).

**Figure 2.**
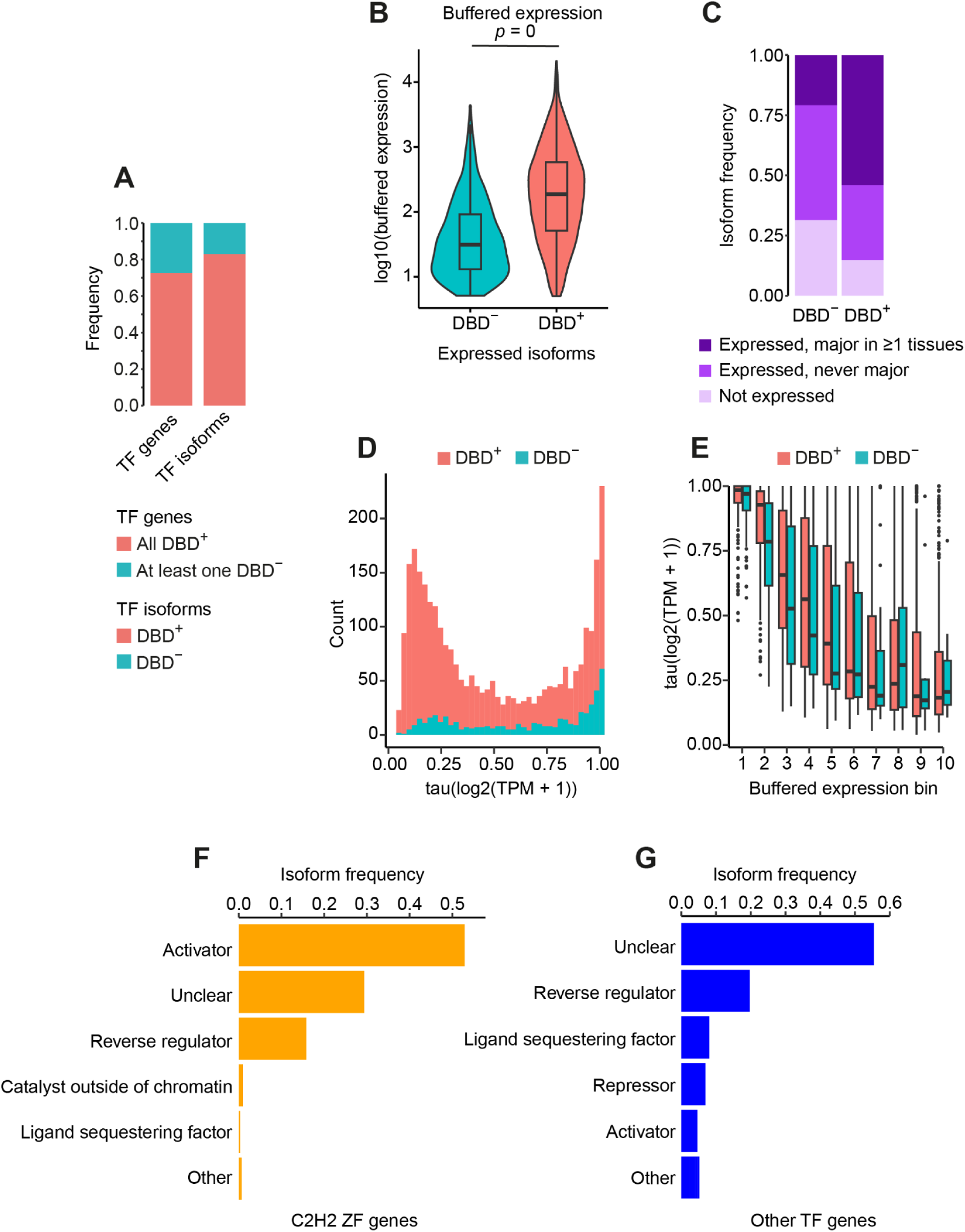
DBD^−^ isoforms of human TF genes are abundant and can be assigned to predicted functional classes. **(A)** The proportion of TF genes with at least one annotated DBD^−^ isoform (left bar) and the proportion of DBD^−^ isoforms among all annotated human TF isoforms (right bar). **(B)** Buffered expression levels of DBD^+^ (*N* = 2,918) and DBD^−^ (*N* = 483) isoforms expressed in adult human tissues (two-sided Wilcoxon rank sum test with continuity correction). The buffered expression level of an isoform is the mean of the top 20% of its expression levels across tissues where the isoform is expressed. The isoform expression levels were normalised across tissue samples using DESeq2 (84). **(C)** Proportions of all annotated DBD^+^ (*N* = 3,427) and DBD^−^ (*N* = 705) isoforms that are not expressed in adult human tissues, expressed in at least one tissue but never become highest-expressed (major), or become major in at least one tissue. **(D)** Distributions of the tissue specificity scores of expressed DBD^+^ and DBD^−^ isoforms. **(E)** Tissue specificity of expressed DBD^+^ and DBD^−^ isoforms, grouped by comparable levels of buffered expression. **(F, G)** Proportions of expressed DBD^−^ isoforms per predicted functional class. The isoforms are stratified into those produced by C2H2 ZF genes (*N* = 310 isoforms) **(F)** and those produced by other TF genes (*N* = 173 isoforms) **(G)**.

In our set of TF genes with at least one DBD^−^ isoform, we found four out of five genes that produce experimentally characterised DBD^−^ isoforms and are included in our compendium: *IKZF1*, *HOXA1*, *TEAD4* and *ZGPAT* **(Figure 1B-C)**. The lack of a DBD^−^ isoform for *IKZF2* **(Figure 1B)** may be due to a removal of this isoform, based on our annotation quality criteria (see **Material and Methods**), if the isoform was present in the full Ensembl annotation. To validate our set of DBD^−^ isoforms at a greater scale, we surveyed publications about human TF isoforms **(Table S2)**, spanning the last three decades, and obtained 26 TF genes with experimentally detected DBD^−^ isoforms **(Table S4)**. We found that our set of 415 TF genes with at least one DBD^−^ isoform was significantly overrepresented among the obtained 26 genes (observed proportion of 0.5, or 13 out of 26 genes, versus the expected proportion of 0.275, or 415 genes with at least one DBD^−^ isoform out of all 1,512 TF genes; *p* = 0.004, hypergeometric test). This fact verifies the biological relevance of our DBD^−^ isoform set.

Next, we determined how the 705 DBD^−^ isoforms that we found in the human genome annotation are distributed across structural families of TF genes. We found that the DBD loss occurs, with different frequencies, in the majority of the families (46 out of 66; **Figure S2**). Namely, the proportion of TF genes with at least one annotated DBD^−^ isoform ranged from 0 to 1, depending on the family, with the two extreme values found almost exclusively in the smallest families **(Figure S2)**. Hence, according to the whole-genome annotation, the loss of a DBD is widespread among human TF genes from various structural families.

As DBD^−^ isoforms are widely annotated across TF genes, we assessed the expression of these isoforms. We chose 43 well-sampled tissues profiled in Genotype-Tissue Expression (GTEx) v8 (83), including 11 brain subtissues, reproductive organs (testis, ovary, uterus and vagina) and other non-brain tissues. We randomly selected 50 samples per tissue to estimate the median expression of each isoform produced by each TF gene in each tissue. We operationally postulated that an isoform is expressed in a tissue if the isoform’s DESeq2-normalised expression level in that tissue is at least 5 (84). In this way, we obtained 3,401 isoforms expressed in at least one tissue. These isoforms were produced by 1,476 TF genes. When we stratified the expressed isoforms into DBD^+^ and DBD^−^, we found that although the DBD^−^ isoforms are expressed lower than the DBD^+^ ones **(Figure 2B)**, 69% (483/705) of all annotated DBD^−^ isoforms are expressed in at least one tissue **(Figure 2C)**. Moreover, using the GTEx-derived transcripts per kilobase per million (TPM) expression values of the expressed isoforms, we additionally selected isoforms that become highest-expressed (major) in at least one tissue. We obtained a set of 2,000 such isoforms, and by intersecting this set with the full set of annotated DBD^−^ isoforms, we found that 21% (147/705) of annotated DBD^−^ isoforms become major in at least one tissue **(Figure 2C)**. Thus, the DBD^−^ isoforms of human TF genes are not only well-represented among all annotated isoforms of these genes, but are also expressed across adult human tissues, with one-fifth of the annotated DBD^−^ isoforms becoming major in particular tissues.

To assess the potential functional importance of expressed DBD^−^ isoforms (*N* = 483), we quantified and compared the tissue specificity of DBD^−^ and DBD^+^ isoforms across the 43 adult human tissues selected before. We calculated isoform tissue specificity as the tau index (85) based on log-scaled isoform TPMs available in GTEx v8. The tau index robustly quantified the tissue specificity of known broadly expressed **(Figure S3A-B)** and tissue-specific **(Figure S3C)** TF genes. We found a pronounced bimodal distribution of the tissue specificity scores of DBD^+^ isoforms **(Figure 2D)**, which suggests that isoforms acting as TFs tend to be expressed either across the majority of adult human tissues or in small tissue subsets. In contrast, the distribution of the tissue specificity scores of DBD^−^ isoforms was biased towards the highest specificity **(Figure 2D)**. However, the lower expression levels of DBD^−^ isoforms may lead to artificially high tissue specificity estimations (see **Material and Methods** for the definition of the tau index). Indeed, **Figure S3D** shows a negative correlation between isoform expression and tissue specificity. Hence, we binned expressed isoforms by their expression level and compared the tissue specificity of DBD^+^ and DBD^−^ isoforms within each bin **(Figure 2E)**. We found that among the lower-expressed isoforms, from bins 2-5, DBD^−^ isoforms are less tissue-specific than DBD^+^ ones, while the higher-expressed DBD^−^ and DBD^+^ isoforms, from bins 6-10, are comparably broadly expressed across tissues. While the lower tissue specificity of the low-expressed DBD^−^ isoforms may suggest their production as "transcriptional noise" (if untranslated), it is also compatible with the possibility that a part of these isoforms are functional, as the DBD^−^ isoforms from bins 2-4 still demonstrate mid to high tissue specificity. On the other hand, the existence of DBD^−^ isoforms produced across the majority of tissues and highly expressed in at least some of them (bins 7-10) is intriguing and suggests their functional relevance.

In sum, the wide distribution of annotated DBD^−^ isoforms across the structural families of TF genes and the expression and tissue specificity of these isoforms in adult human tissues suggest that DBD^−^ isoforms may have functional roles.

### Non-DNA-binding domains predict functional classes of DBD^−^ isoforms

To predict general functional roles (functional classes) of expressed DBD^−^ isoforms, we used domain descriptions from the InterPro database (89) and extensively searched literature to functionally annotate 262 unique types of non-DNA-binding domains (non-DBDs) matched in annotated human TF isoforms **(Table S5)**. We subset this non-DBD annotation for expressed DBD^−^ isoforms, stratifying the isoforms into those produced by Cys_2_His_2_ (C2H2) zinc finger (ZF) genes (*N* = 310 DBD^−^ isoforms) and by other TF genes (*N* = 173 DBD^−^ isoforms). We found that 71% (220/310) of C2H2 ZF DBD^−^ isoforms and 50% (86/173) of other DBD^−^ isoforms contain non-DBDs. As expected, the most frequent functional types of non-DBDs in the C2H2 ZF DBD^−^ isoforms were a repressor domain (KRAB), an oligomerization domain (SCAN) and a dimerization domain (BTB/POZ) **(Figure S3E)**. The DBD^−^ isoforms produced by other TF genes contained a wider range of non-DBDs, including various dimerization, ligand-binding, transcription regulation and other types of domains **(Figure S3F)**. Consequently, these domains may confer function to DBD^−^ isoforms and could be used to predict functional classes of these isoforms. However, 29% (91/310) of DBD^−^ isoforms produced by C2H2 ZF genes and 50% (87/173) of other DBD^−^ isoforms do not have any domains, which precluded their functional characterization.

Next, using the obtained functional non-DBD annotation **(Table S5)** and previously studied examples of functional DBD^−^ isoforms **(Figure 1B-C)**, we assigned functional classes to all DBD^−^ isoforms with at least one non-DBD, stratifying the isoforms into those produced by C2H2 ZF genes and by other TF genes **(Figure 2F-G; Table S6-S7)**. Sixty-nine percent (213/310) of DBD^−^ isoforms produced by C2H2 ZF genes were predicted to be activators or "reverse regulators" (regulators of transcription whose effect is opposite to the one exerted by a DBD^+^ isoform with the same non-DBDs; **Figure 2F**). This prediction is based on the presence of repressor domains or less specifically defined transcription regulation domains which, according to experimentally studied cases **(Figure 1C)**, could sequester transcriptional corepressors (in general, transcriptional cofactors) from the chromatin and thus block the regulation that the cofactors would otherwise exert. Thirty-nine percent (68/173) of DBD^−^ isoforms expressed by other TF genes were predicted to be reverse regulators, ligand sequesterers or transcriptional regulators (activators or repressors; **Figure 2G**). Overall, 71% (219/310) of DBD^−^ isoforms expressed by C2H2 ZF genes and 45% (77/173) of DBD^−^ isoforms expressed by other TF genes were assigned a predicted functional class **(Table S7)**, suggesting the importance of further experimental efforts to understand the possible roles of DBD^−^ isoforms. Importantly though, the rest of the DBD^−^ isoforms were not assigned a functional class mainly because of the lack of non-DBDs (the "Unclear" class in **Figure 2F-G**; also see **Figure S3G-H**). This fact highlights the need to study intrinsically disordered regions to fully understand the possible roles of these domainless isoforms.

In total, at least 71% of DBD^−^ isoforms expressed by C2H2 ZF genes and 45% of DBD^−^ isoforms expressed by other genes have a potential for a functional role, based on non-DBDs. This suggestion is additionally supported by the existence of highly tissue-specific DBD^−^ isoforms expressed on par with some DBD^+^ isoforms, as well as the presence of tissue-non-specific DBD^−^ isoforms highly expressed in at least a subset of adult human tissues.

### Examples of highly expressed tissue-non-specific DBD^−^ isoforms

The highly expressed tissue-non-specific DBD^−^ isoforms that we found above (**Figure 2E**, expression bins 7-10) may functionally interact with TFs, including the DBD^+^ isoforms of the same TF genes, or play their own roles. To reveal the expression profiles and functional classes of such DBD^−^ isoforms, we selected them from the expression bins 7-10 using the maximum tau index cutoff of 0.25. We obtained 33 DBD^−^ isoforms **(Table S8)** and inspected them manually. We found that these DBD^−^ isoforms are often coexpressed in the same tissue with the DBD^+^ isoforms of the same TF gene. Below, we describe particular examples of such DBD^−^ isoforms that, to the best of our knowledge, have not been functionally characterised in adult human tissues.

The highly expressed DBD^−^ isoform of a C2H2 ZF gene *ZNF197* is tightly coexpressed with a DBD^+^ isoform of the same gene across adult human tissues **(Figure 3A)**. Both isoforms have a transcription repression KRAB domain and an oligomerization SCAN domain. Although, according to our classification, this DBD^−^ isoform could activate transcription through the sequestration of corepressors and the DBD^+^ isoform from the chromatin **(Figure 1B-C)**, Li and colleagues have shown that this DBD^−^ isoform, which they called VHLaK, facilitates transcription repression via binding the VHL protein and inhibiting the transcriptional activity of the HIF1A TF (90). Therefore, the broad expression profile of the DBD^−^ isoform in **Figure 3A** suggests that the transcription repression mediated by this isoform and described by Li and colleagues in cell lines may take place across a wide range of adult human tissues, depending on the expression levels of HIF1A and VHL. However, VHLaK may still compete for corepressors with the DBD^+^ isoform of *ZNF197* or with other KRAB domain-containing transcriptional repressors, if they are coexpressed in the same cells **(Figure 1C)**. This competition may lead to the re-balancing of direct (via the DBD^+^ isoform) and indirect (via the DBD^−^ isoform) transcription repression, depending on the expression levels of the two *ZNF197* isoforms and their binding partners, which calls for a future experimental investigation.

**Figure 3.**
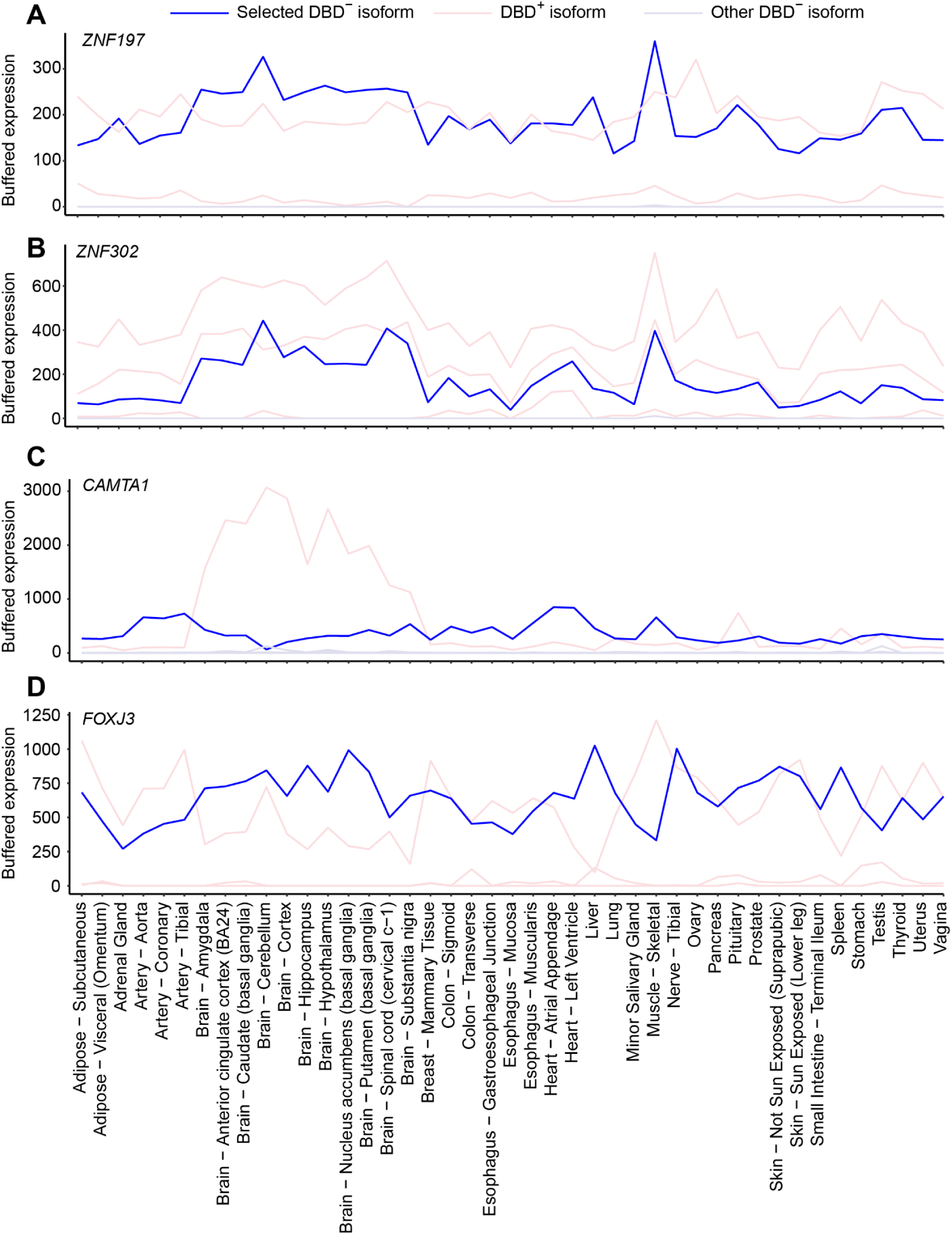
Examples of highly expressed tissue-non-specific DBD^−^ isoforms ("Selected DBD^−^ isoform") from expression bins 7-10, Figure 2E, produced by *ZNF197* (isoform ENST00000383745, **A**), *ZNF302* (isoform ENST00000505365, **B**), *CAMTA1* (isoform ENST00000473578, **C**) and *FOXJ3* (isoform ENST00000372571, **D**). The buffered expression level of an isoform is the mean of the top 20% of its expression levels across tissues where the isoform is expressed. The isoform expression levels were normalised across tissues with DESeq2.

Another C2H2 ZF gene, *ZNF302*, coexpresses its highly expressed tissue-non-specific DBD^−^ isoform with two DBD^+^ isoforms, and the expression profiles of the three isoforms are well-coordinated **(Figure 3B)**. Similarly to the previous example, all the three isoforms have a KRAB domain. Therefore, the DBD^−^ isoform of *ZNF302* could repress transcription by inhibiting transcriptional activity of TFs (similar to VHLaK) or attenuate transcription repression by competing for corepressors with the DBD^+^ isoforms of *ZNF302* and other repressors **(Figure 1C)**. However, the role of the DBD^−^ isoform has not been characterised, and the functional role of the gene is still poorly understood. Both facts suggest the importance of experimental research into the isoforms of *ZNF302*.

*CAMTA1*, a TF gene with a CG-1 type of the DBD, has a highly and broadly expressed uncharacterized DBD^−^ isoform which cannot be assigned to a functional class due to a lack of domains. However, if intrinsically disordered, it may still bind DNA, RNA or proteins, which could be validated experimentally. *CAMTA1* expresses the DBD^−^ isoform across the majority of adult human tissues, where it also expresses a DBD^+^ isoform **(Figure 3C)**. However, the DBD^+^ isoform clearly dominates in the brain, which corresponds to previously studied roles of this TF in episodic memory (91), Purkinje cell function and survival (92) and in glioblastoma suppression (93). Additionally, in arteries, heart and skeletal muscles the DBD^−^ isoform is expressed considerably higher than the DBD^+^ isoform. Therefore, *CAMTA1* effectively switches a DBD^+^ (potential TF) isoform dominating in the brain for a domainless isoform dominating in particular non-brain tissues.

Finally, *FOXJ3* demonstrates an anticorrelated expression of a DBD^+^ and a DBD^−^ isoform across adult human tissues **(Figure 3D)**. As the DBD^−^ isoform lacks domains, it cannot be assigned a functional class. However, similarly to the previous example, *FOXJ3* may effectively switch a DBD^+^ (potential TF) isoform for a DBD^−^ isoform between various tissues. Indeed, upon visual inspection, the DBD^+^ isoform is expressed considerably higher, than the DBD^−^ isoform, in the adipose tissue, arteries, uterus and testis, but most clearly in skeletal muscles where the FOXJ3 TF was experimentally suggested to regulate the fibre cell identity and regeneration (94). In contrast, the DBD^−^ isoform dominates in particular brain tissues and the pituitary gland, as well as in the liver and spleen. Consequently, we expect the TF function of *FOXJ3* to be considerably attenuated in these tissues.

In total, the highly and broadly expressed DBD^−^ isoforms may affect transcription independently or through interactions with the DBD^+^ isoforms of the same or other TF genes. These options highlight the need for experimental investigations of the possible roles of these and other DBD^−^ isoforms, still uncharacterized in adult human tissues. Intriguingly, the isoform expression profiles shown above reveal putative switching patterns between the highest expressed (major) isoforms **(Figure 3A, C-D)**. Therefore, we decided to study such major isoform switches systematically and at a genome-wide scale.

### Human TF genes switch major isoforms between adult human tissues

Isoform switching plays a prominent role in meiosis (95), cell differentiation (18), development (96) and disease (97–100). To reveal potential switches of the 2,000 major isoforms of human TF genes across the 43 selected adult human tissues, we started by formally defining a switch. We postulated that a switch between major isoforms *K* and *J* of the same TF gene is an exchange of the isoform *K* expressed as major in one set of tissues for the isoform *J* expressed as major in another set of tissues, such that the difference between the expression levels of the two isoforms in any tissue from the two sets is at least 20% **(Figure 4A)**. After considering statistical tests to define a switch, we instead preferred the definition based on a relative difference in isoform expression. We reasoned that this relative difference has to be considerable to suggest a biological effect but does not have to be significantly greater than the most part of differences in other pairs of major isoforms. Additionally, we are interested in switches occurring between any subsets of tissues, which would have entailed hundreds of pairwise tests for a switch between all pairs of tissues. Finally, we are not testing the significance of the sizes of tissue sets taking part in a switch, because a switch between any number of tissues of any kind could be functionally relevant.

**Figure 4.**
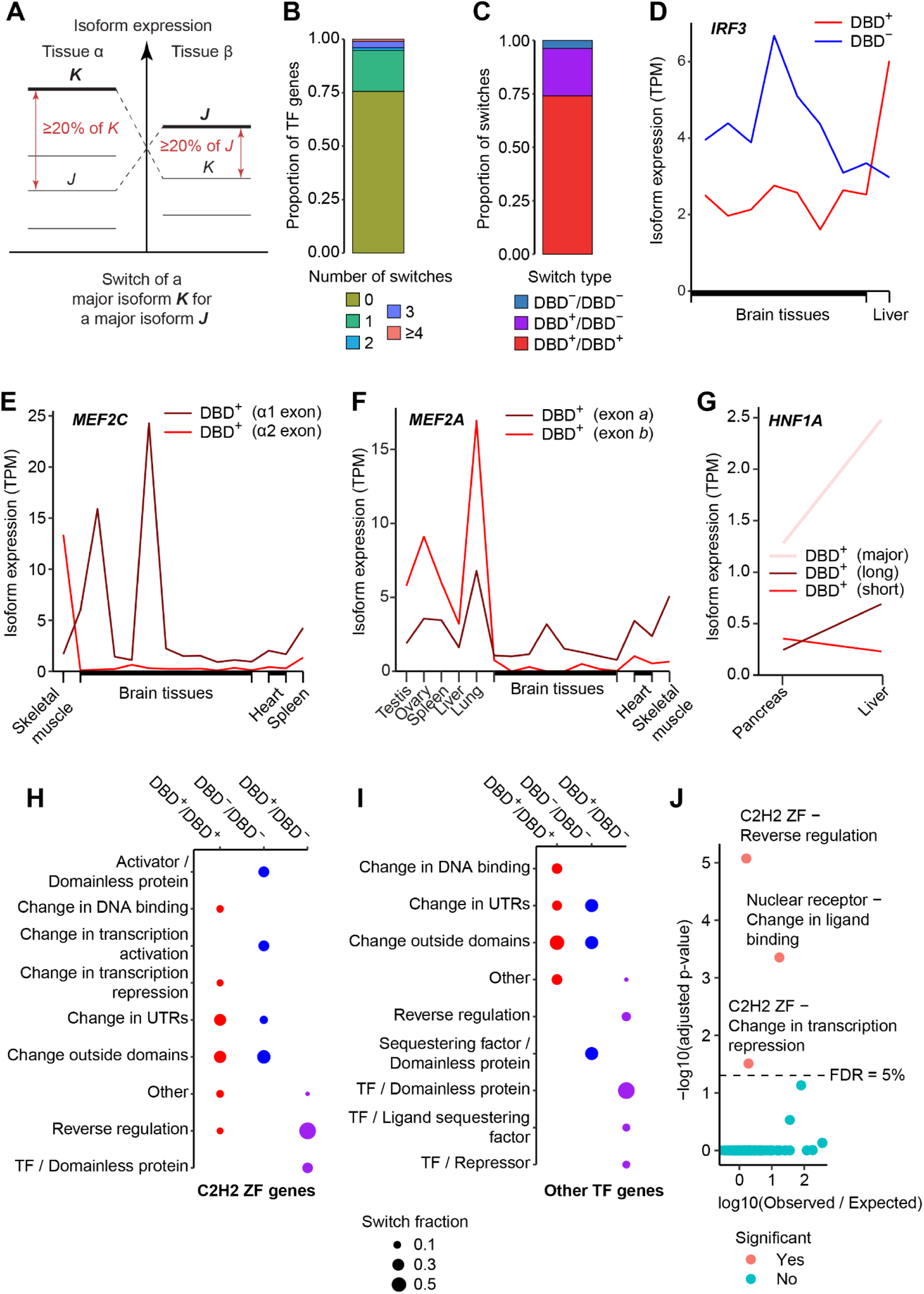
Human TF genes switch major isoforms across adult human tissues with predictable functional consequences. **(A)** Switch of a major isoform *K* for a major isoform *J* of the same TF gene between tissues **α** and **β**. Switching major isoforms can be coexpressed with other isoforms of the same TF gene in the same tissues. **(B)** Proportions of human TF genes by the number of produced switches. **(C)** Proportions of switches by type (DBD^+^/DBD^+^, DBD^+^/DBD^−^ and DBD^−^/DBD^−^). **(D-G)** Switches of the major isoforms of TF genes that were previously validated experimentally and that are closely recapitulated in our analysis: *IRF3* **(D)**, *MEF2C* **(E)**, *MEF2A* **(F)**, *HNF1A* **(G)**. **(H-I)** Predicted functional classes of switches produced by C2H2 ZF genes **(H)** or other TF genes **(I)**. "Other" switch classes in each panel summarise classes containing less than 5% of switches. **(J)** TF gene families significantly overrepresented in particular switch classes (FDR = 5%). We performed pairwise hypergeometric tests of the overrepresentation of TF gene families (*N* = 66) in switch classes (*N* = 36) and corrected raw p-values using the Benjamini-Hochberg procedure. We obtained the following significant overrepresentations: C2H2 ZF genes in the switch class "Reverse regulation" (adjusted *p* = 8.5e-06), nuclear receptors in the switch class "Change in ligand binding" (adjusted *p* = 4.4e-04) and C2H2 ZF genes in the switch class "Change in transcription repression" (adjusted *p* = 3.1e-02).

According to the proposed definition, we obtained 536 switches between 816 major isoforms expressed by 360 TF genes **(Table S9)**. Therefore, 24% (360/1,476) of expressed TF genes produced at least one switch, and we found that the majority of these genes produced only one switch **(Figure 4B)**. Moreover, the vast majority (74%) of all switches occurred between two DBD^+^ isoforms (we denote these switches DBD^+^/DBD^+^), while only 22% of switches were DBD^+^/DBD^−^ **(Figure 4C)**. This is expected, given the rarity of the DBD^−^ isoforms in comparison to the DBD^+^ ones. Finally, the DBD^−^/DBD^−^ switches comprised the remaining 4% of major isoform switches, supporting the possibility of a functional role for at least some DBD^−^ isoforms **(Figure 4C)**.

To confirm the reliability of our switch calling method, we checked the presence in our switch list of any known isoform switches produced by human TF genes or orthologous TF genes in other mammalian species across normal adult tissues. Although, based on literature, the vast majority of TF isoform switches were studied in development or cancer, we found 8 TF genes producing experimentally verified major isoform switches between tissues present in our tissue set: *HNF1A* (101, 102), *HNF4A* (103), *IRF3* (104), *MEF2A* and *MEF2D* (105), *MEF2C* (106), *TCF4* (107, 108) and *TEAD1* (109). We exactly reproduce switches by *IRF3* **(Figure 4D)** and *MEF2C* **(Figure 4E)**, matching tissues taking part in the switches and the domain structures of switching major isoforms. We also closely recapitulate the switch by *MEF2A* **(Figure 4F)**, identifying the same switching tissues; however, our switch likely reflects the differential inclusion of a different exon, due to mismatches between the Ensembl exon annotation of this gene and the annotation shown in the article by Ab Hakim et al. Finally, we identified the switch produced by *HNF1A*, although the switching isoforms in our dataset are not expressed as major **(Figure 4G)**. This is likely due to differences in the isoform quantification, or the sets of detected isoforms, between the GTEx database and the two studies by Harries et al. In total, the close reproduction of the four known major isoform switches supports the genome-wide application of our switch calling method based on the GTEx data.

Noteworthy, we did not recapitulate experimentally detected switches produced by *HNF4A*, *MEF2D*, *TCF4* and *TEAD1*. For *HNF4A* and *MEF2D*, we detect switches of the expected DBD^+^/DBD^+^ type **(Table S9)** but their tissue sets do not match those listed by Nakhei et al. and Zhu et al. Also, for *TCF4*, we detect a switch between the expected tissues (testis and lung), but the switch is DBD^+^/DBD^−^ instead of the expected DBD^+^/DBD^+^. Finally, for *TEAD1* we do not find switches, as, according to our data, it has the same major isoform across all tissues, and the lower-expressed isoforms do not switch. In sum, these discrepancies suggest a difference in the isoform expression quantification or the sets of quantified isoforms between the GTEx database and the corresponding studies.

### Domains of switching isoforms suggest functional outcomes of switches

After obtaining the genome-wide set of major isoform switches (*N* = 536) and establishing the presence of several experimentally validated switches, we set out to predict possible functional outcomes of the switches we found. We started by testing the overrepresentation of Gene Ontology (GO) Biological Process, Molecular Function and Cellular Component terms (110, 111), as well as the Molecular Signatures Database (MSigDB) Hallmark and C2 Canonical Pathways (112–114), in switching TF genes. As a background, we used all TF genes expressing more than one major isoform. With this, we did not find any GO term overrepresentation under a false discovery rate (FDR) of 10%, which suggested that the major isoform switches are not constrained to particular molecular pathways. Furthermore, we tested possible overrepresentation of TF gene families in the switching TF genes and again did not find significant results under FDR = 10%. This fact additionally suggests the independence of major isoform switching from the structural class of a DBD encoded by a TF gene. Finally, the three types of switches (DBD^+^/DBD^+^, DBD^+^/DBD^−^ and DBD^−^/DBD^−^) also did not demonstrate significant overrepresentation of GO terms, pathways from the hallmark or canonical sets, or TF gene families. Overall, the observed lack of significant GO term overrepresentations suggests that major isoform switches are non-preferentially distributed across molecular processes and structural TF gene families.

Next, based on domains present in switching isoforms and on examples of validated functional differences between alternative isoforms of the same TF gene **(Figure 1, S1)**, we predicted functional classes of major isoform switches **(Figure 4H-I; Table S9)**. Because switches produced by C2H2 ZF genes (*N* = 275 switches) comprise 51% of all major isoform switches (*N* = 536), we predicted functional classes separately for C2H2 ZF switches **(Figure 4H)** and for other switches **(Figure 4I)**. Strikingly, we found that the majority of DBD^+^/DBD^+^ switches from both switch groups (63% of C2H2 ZF DBD^+^/DBD^+^ switches and 61% of other DBD^+^/DBD^+^ switches) do not affect domains and, therefore, do not allow us to assign a functional class based on domain differences. Instead, these switches either changed a protein sequence outside domains or changed untranslated regions (UTRs) **(Figure 4H-I)**.

One of the well-represented functional classes of DBD^+^/DBD^+^ switches in both groups of TF genes is a "Change in DNA binding" (10% of C2H2 ZF DBD^+^/DBD^+^ switches and 19% of other DBD^+^/DBD^+^ switches) **(Figure 4H-I)**. In C2H2 ZF switches, it is exclusively explained by changing numbers of zinc fingers **(Figure 1A, *left*)**, while in other switches it is inferred from changes in the DBD sequence **(Figure 1A, *middle* and *right*)**. These switches may modulate the target-specificity of a TF **(Figure 1A)**, hence differentially regulating the corresponding molecular pathways. Otherwise, although in these switches both major isoforms contain a DBD, the switches may still replace a DNA-binding major isoform with a non-DNA-binding one if the number or configuration of zinc fingers is not sufficient for DNA binding or if the sequence of a non-ZF DBD becomes too short or lacks a critical part.

Other functional classes of DBD^+^/DBD^+^ switches produced by C2H2 ZF genes include "Change in transcription repression" (due to changes in the KRAB domain sequence, 9%) and "Reverse regulation" (inferred from the differential inclusion of the KRAB domain, 8%) **(Figure 4H; Table S9)**. These switches suggest that the expression of different major isoforms by the KRAB-encoding C2H2 ZF genes may modulate the strength or specificity of transcription repression (**Figure 1G** and, by extrapolation, **Figure 1F**) or even abolish the repression **(Figure 1E)**.

The vast majority of DBD^+^/DBD^−^ switches produced by C2H2 ZF genes (*N* = 76 switches) are predicted to reverse the transcription regulation effect (*N* = 55), mainly due to the differential inclusion of the full set of zinc fingers in the presence of the KRAB domain which would repress transcription in a DBD^+^ isoform but likely sequester corepressors in a DBD^−^ isoform **(Figure 1C)**. In other TF genes, DBD^+^/DBD^−^ switches mainly replace a DBD^+^ major isoform with a domainless one, thus precluding domain-based functional predictions. This fact underscores the necessity of studying intrinsically disordered regions of TF gene products to fully understand their function. However, other DBD^+^/DBD^−^ switches of non-C2H2 ZF genes can be assigned to predicted functional classes based on the presumed sequestration of partner TFs via a dimerization domain ("Reverse regulation"; **Figure 1B**), coactivators via a transcription activation domain ("TF / Repressor"; **Figure 1C**) or ligands via a ligand-binding domain ("TF / Ligand sequestering factor", by analogy to the other two cases).

Through the functional classification of major isoform switches, we effectively assigned each of the 360 switching TF genes to one or more classes based on the switches that the gene produces. Therefore, we next asked if switches of particular functional classes tend to be produced by TF genes from particular families. To this end, we tested the overrepresentation of the 66 TF gene families in the 36 functional classes of switches that we defined **(Figure 4J)**. After controlling for multiple testing, we found that C2H2 ZF genes are overrepresented in switch classes "Reverse regulation" and "Change in transcription repression," while nuclear receptors are overrepresented in the switch class "TF / Change in ligand binding" defined by the changing sequence of the ligand-binding domain **(Figure 4J)**. Therefore, the majority of switch classes are produced by TFs from different families, while the three switch classes listed above are produced mainly by C2H2 ZF or nuclear receptor genes.

In total, our set of 536 predicted major isoform switches drastically expands the collection of several experimentally validated TF isoform switches between adult human tissues described in the previous section. The fact that almost a quarter of all human TF genes that are expressed in adult human tissues produce at least one major isoform switch, as well as our functional predictions, suggest that the switches may represent a widespread, but mostly uncharacterized, layer of regulation of transcription and, potentially, of other processes in the cell.

### Human TF genes switch major isoforms between the brain and non-brain tissues

To reveal the tissue specificity of the major isoform switches, we clustered both the switches and the tissues according to the type of the switching major isoform in each tissue: DBD^+^, or DBD^−^, or “null” if the switching TF gene is not expressed in the tissue or if its major isoform expressed in the tissue does not take part in a switch **(Figure 5)**. We used complete linkage (CL) clustering to highlight the DBD^+^/DBD^−^ switches **(Figure 5A-B, S4)** and Ward’s D2 linkage (WD2L) clustering to highlight the DBD^+^/DBD^+^ switches **(Figure 5A, 5C, S5)**. We used two different linkages, as a visual inspection has shown that they lead to more consistent clusters of DBD^+^/DBD^−^ or DBD^+^/DBD^+^ switches, respectively. We found that brain tissues cluster separately from non-brain tissues in both cases, suggesting a considerable proportion of brain-to-non-brain switches among all the switches we found. Indeed, the CL cluster 5 demonstrates a striking brain-to-non-brain expression pattern **(Figure 5B)**, with the corresponding TF genes producing major DBD^+^ isoforms outside the brain and major DBD^−^ isoforms within the brain. Additionally, the WD2L cluster 3 highlights another brain-to-non-brain switch pattern **(Figure 5C)**, with one major DBD^+^ isoform expressed outside the brain and another major DBD^+^ isoform expressed in the brain.

**Figure 5.**
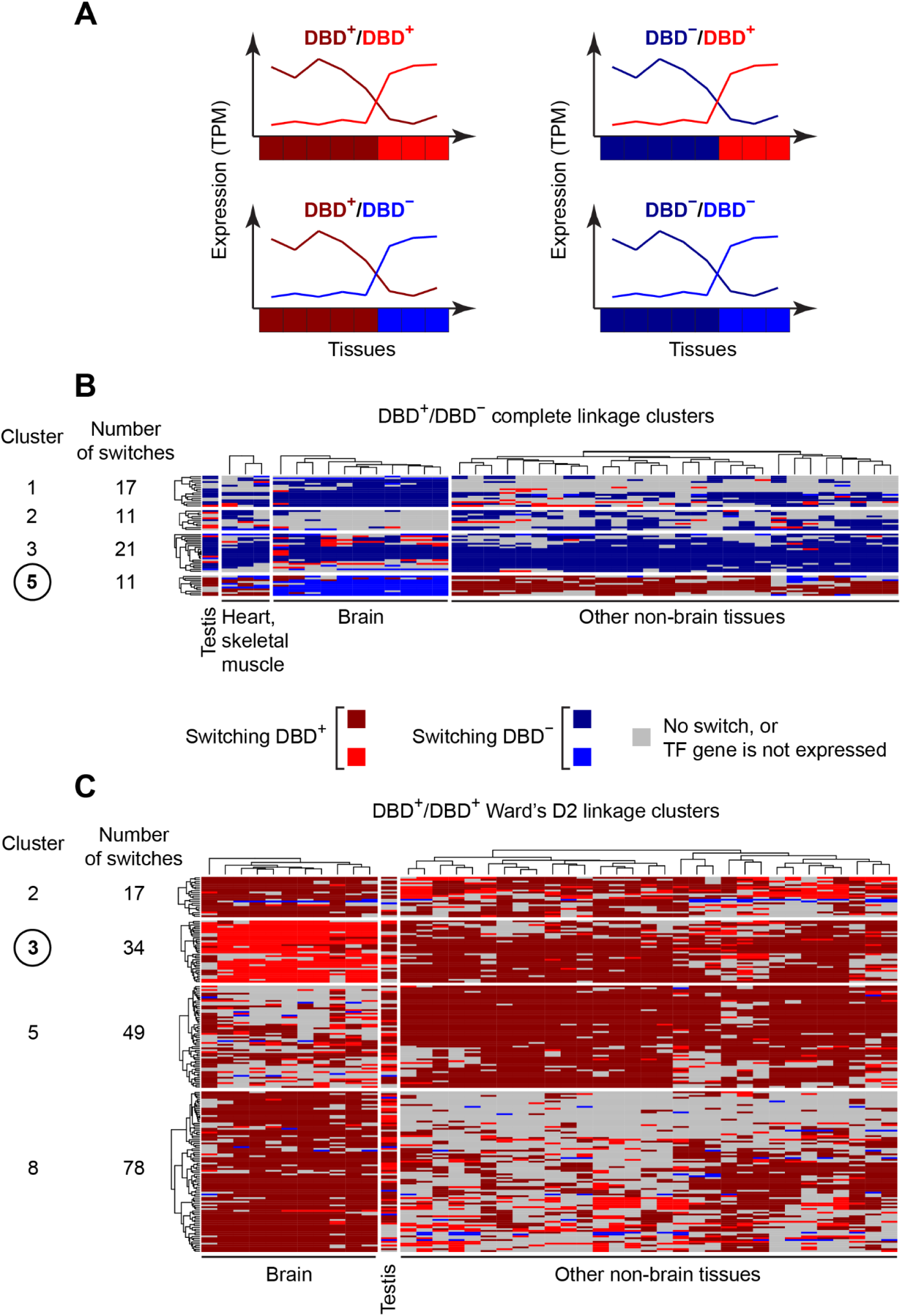
Expression patterns of major isoform switches between the brain and non-brain adult human tissues. **(A)** Schematics of the four possible combinations of switching major DBD^+^ and DBD^−^ isoforms and the corresponding representations of these switches as rows of the heatmaps in panels **B** and **C**. Shades of red denote switching DBD^+^ isoforms; shades of blue denote switching DBD^−^ isoforms; darker shades denote switching isoforms that are major in a higher number of tissues than their counterparts. If switching isoforms are major in an equal number of tissues, then their colours are assigned randomly. **(B-C)** Switches (in rows) and tissues (in columns) are hierarchically clustered based on the types of switching major isoforms (DBD^+^ or DBD^−^) and the tissue-specificity of their expression. See **Material and Methods** for details. **(B)** Selected clusters of DBD^+^/DBD^−^ switches from the hierarchical clustering with the complete linkage (see **Figure S4** for the full set of clusters). **(C)** Selected clusters of DBD^+^/DBD^+^ switches from the hierarchical clustering with Ward’s D2 linkage (see **Figure S5** for the full set of clusters).

The rest of the clusters shown in **Figure 5B-C** represent more tissue-specific switch patterns. Thus, in the CL clusters 1, 2 and 3, TF genes do not produce TFs (or produce them at lower levels) across the majority of tissues where the genes are expressed, but switch a non-TF product for a TF (encoded by a major DBD^+^ isoform) in particular tissues within the brain (cluster 3) or outside the brain (clusters 1 and 2). Similarly, in the WD2L clusters 2, 5 and 8, a broadly expressed DBD^+^ (potential TF) isoform switches for another DBD^+^ isoform, or even a rare DBD^−^ isoform, in particular tissues, mostly within the brain (cluster 5) or outside the brain (clusters 2 and 8). Overall, the 8 selected clusters suggest extensive major isoform switches of human TF genes between the brain and the non-brain tissues, leading us to an investigation of possible functional outcomes of these switches.

To obtain functional insights into the brain-to-non-brain switches within each selected cluster, we tested the overrepresentation of GO terms, the hallmark and the canonical pathways from MSigDB, as well as TF gene families, in the sets of TF genes that produce switches in each cluster. In brief, this test did not show significant overrepresentations (FDR = 10%), suggesting that the observed brain-to-non-brain switch patterns occur across various molecular processes and TF gene families.

As the overrepresentation analysis did not yield significant results, we selected only the two clusters with the clearest brain-to-non-brain expression patterns (the CL cluster 5 and the WD2L cluster 3) and manually checked the TF genes that produce them for recurring physiological processes in which they may participate. Among the 11 TF genes from the CL cluster 5, only 5 genes (*IRF6*, *PRDM2*, *RARA*, *THAP4* and *ZNF568*) had predicted or established physiological roles, and among these, four genes took part in various developmental processes: *IRF6* – in epidermal development (115); *PRDM2*, as a *PRDM* factor, likely, – in neural differentiation (116); *RARA* – in spermatogenesis (117, 118) and *ZNF568* – in placenta morphogenesis (119) and maintenance of neural stem cells (120). However, the relevance of these developmental processes to the brain-to-non-brain isoform switches in adult tissues is unclear.

Among the 34 TF genes from the WD2L cluster 3, 27 genes had predicted or established physiological roles. We found two broad physiological processes recurring in the characteristics of these 27 genes. The first was the central nervous system (CNS) development and neuroprotection represented by genes *ADNP*, *BCL11A*, *LHX6*, *MEF2A*, *MEIS1*, *ZNF592* and *ZBTB45*, while the second broadly defined process was lipid metabolism and adipocyte differentiation represented by genes *JAZF1*, *PPARG*, *SREBF1*, *TFEB* and *ZNF202* (see **Table S10** for references). While switches between major isoforms of the first group of genes may be directly related to their CNS-specific functions, the switches produced by the second group of genes suggest differences in lipid metabolism within and outside the brain.

### Pronounced brain-to-non-brain major isoform switches suggest functional effects

Above, we uncovered intriguing brain-to-non-brain switches between DBD^−^ and DBD^+^ major isoforms (CL cluster 5; **Figure 5B**) and between DBD^+^ major isoforms (WD2L cluster 3; **Figure 5C**). Based on the domain sets of switching isoforms and on available literature, we set out to manually characterise these 45 switches to obtain a detailed view of their putative functional consequences. Here, we describe and interpret a selected subset of these switches which, to the best of our knowledge, have not been experimentally characterised.

A C2H2 ZF gene *ZNF470* switches a DBD^+^ major isoform, containing a ZF array and a KRAB domain and expressed in non-brain tissues, for a KRAB-only major DBD^−^ isoform expressed in the brain **(Figure 6A, S6A; Table S9)**. Based on our compendium of functionally characterised alternative TF isoforms, we predict that the DBD^−^ major isoform could sequester corepressors from the chromatin in the brain **(Figure 1C)**, in this way attenuating or abolishing the repression of ZNF470 target genes. In contrast, the major DBD^+^ isoform outside the brain could still repress transcription in non-brain tissues. Also, another three C2H2 ZF genes, *ZNF530*, *ZNF571* and *ZNF208*, produce DBD^+^/DBD^−^ brain-to-non-brain switches with the same predicted functional consequences **(Figure S6B-D)**. Importantly, all the four TF genes are poorly characterised, and nothing is known about the functions of their isoforms. Our results suggest the necessity of experimental studies of these genes, especially in brain cell types.

**Figure 6.**
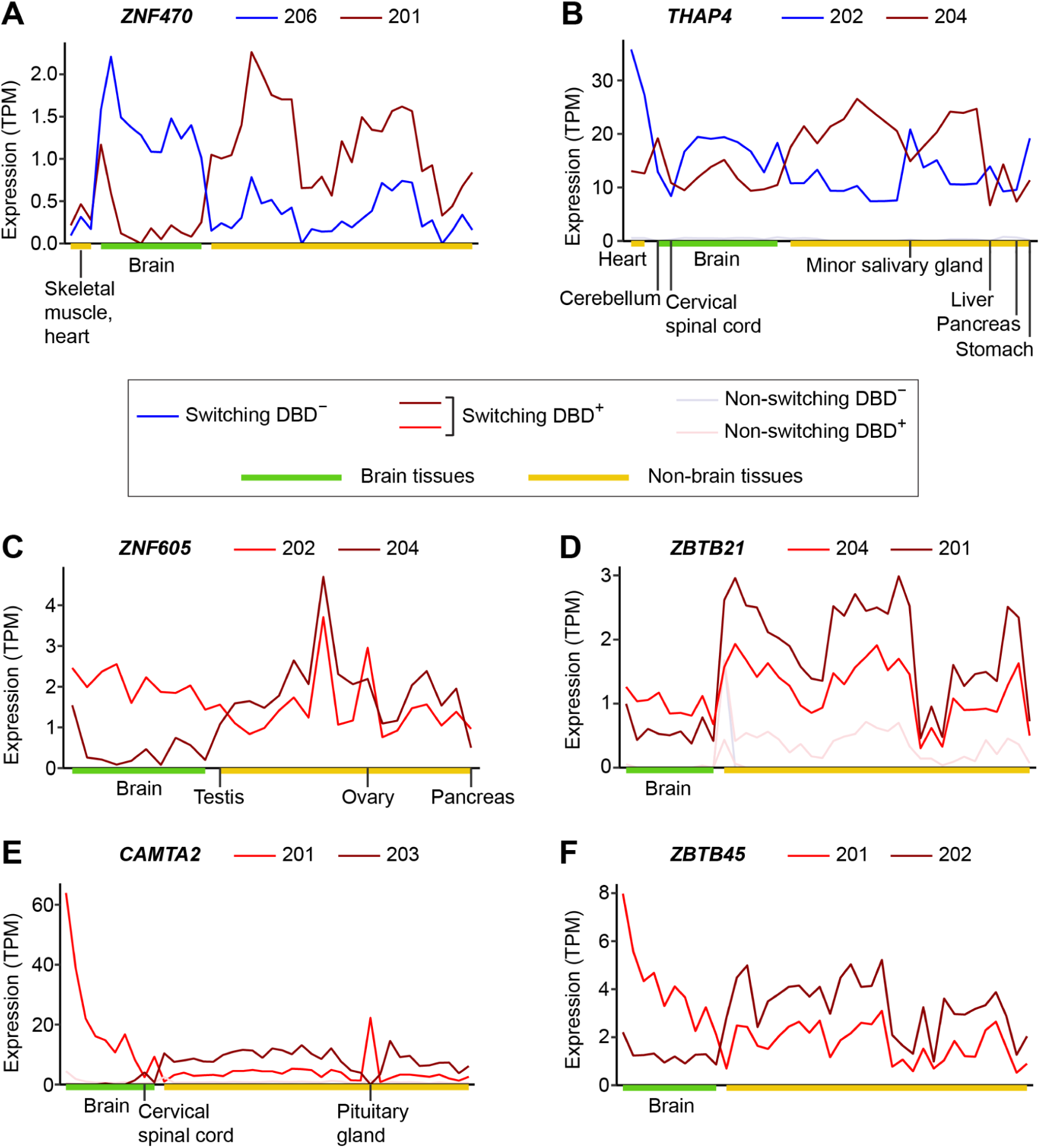
Selected major isoform switches between the brain and non-brain tissues from the complete linkage switch cluster 5 (DBD^+^/DBD^−^ switches in panels A and B) and the Ward’s D2 linkage cluster 3 (DBD^+^/DBD^+^ switches in panels C-F). See Figure 5A for the description of the colour scheme for switching isoforms. The tissue order along the x axis in the panels **A** and **B** corresponds to the tissue order in Figure 5B and **Figure S4**, while the tissue order in the panels **C-F** corresponds to the one in Figure 5B and **Figure S5**. Only the tissues where the switch occurs are represented, and particular tissues are labelled. The same switches with the full set of tissue labels are shown in **Figure S6A, S7A-B, S8A, S9, S10A**. Switching isoforms are denoted according to their identifiers in Ensembl v99. For instance, panel **A** shows a switch between a major DBD^−^ isoform *ZNF470*-206 and a major DBD^+^ isoform *ZNF470*-201. The switches are produced by *ZNF470* **(A)**, *THAP4* **(B)**, *ZNF605* **(C)**, *ZBTB21* **(D)**, *CAMTA2* **(E)**, *ZBTB45* **(F)**.

In a more complex switch, *THAP4* exchanges a full DBD^+^ major isoform dominating in many non-brain tissues, as well as in the cervical spinal cord and cerebellum, for a DBD^−^ major isoform dominating in the rest of the brain tissues and in particular non-brain tissues **(Figure 6B, S7A; Table S9)**. THAP4 binds heme and nitrogen oxide and converts peroxynitrite into nitrate (121–123), although the effect of these activities on the protein’s role in transcription regulation is still unclear. On the other hand, according to our analysis, both switching isoforms have a heme-binding domain. Therefore, we predict that the brain-dominating DBD^−^ isoform may serve as a heme sequestering factor, either altering the regulatory or catalytic effect of the DBD^+^ isoform (if the two isoforms are coexpressed in the same cells) or functioning on its own (in this way, sequestering heme from any other heme-binding molecules).

Among DBD^+^/DBD^+^ major isoform switches, a switch produced by *ZNF605* changes the sequence of a KRAB domain, likely leading to differential transcription regulation of ZNF605 target genes between the brain and non-brain tissues **(Figure 6C, S7B; Table S9)**. The difference in the KRAB domain sequence could manifest in different sets of target genes (**Figure 1F**, by extrapolation) or different transcription repression efficiencies **(Figure 1G)** of the two isoforms. As their relative expression levels are close outside the brain, the two isoforms could produce a combined transcription repression effect in non-brain tissues, if coexpressed in the same cells; however, effectively, only one of the two isoforms is expressed in the majority of the brain tissues, hence defining the brain-specific set of repressed targets. Of note, *ZNF605* may play a role in the hypothalamus development (124), which supports the biological relevance of the described switch.

*ZBTB21*, a transcriptional repressor (125) regulating neurodevelopment (126), produces a DBD^+^/DBD^+^ major isoform switch where a part of a C2H2 ZF array is differentially included in switching isoforms, with the brain-dominating isoform containing fewer zinc fingers **(Figure 6D, S8A; Table S9)**. This alternative splicing event likely leads to different DNA-binding affinities or specificities of the two isoforms and, consequently, to a difference in their cognate sets of target genes **(Figure 1A, left)**. If the two isoforms are coexpressed in the same cells, their combined regulatory effect may depend on their abundance ratio which inverts between the brain and non-brain tissues. Intriguingly, Wang and colleagues recently established ZBTB21 as a facilitator of cohesin-chromatin binding, implicating this factor in the regulation of chromatin conformation (127). In the light of this evidence, the described switch might also differentially affect the 3D genome architecture between the brain and other organs.

Along with *ZBTB21*, the following zinc finger TF genes also change the number of C2H2 ZF domains in their major DBD^+^ isoforms that switch between the brain and non-brain tissues: *ZEB2* **(Figure S8B; Table S9)**, a versatile developmental transcriptional repressor (128) taking part in the development of the nervous system (129–131); *JAZF1* **(Figure S8C; Table S9)**, also a transcriptional repressor and a regulator of lipid and glucose metabolism in the hypothalamus and peripheral tissues (132–134); and *ZNF500* **(Figure S8D; Table S9)**, a zinc finger TF gene with an unknown physiological role. However, Ma and co-authors have found that ZNF500-p53 dimerization increases p53 stability and that the dimerization requires the presence of C2H2 ZF domains in ZNF500 (135). At present, we do not know which C2H2 ZF domain(s) of ZNF500 bind p53, but if one of these domains is the zinc finger included in the brain-dominating major DBD^+^ isoform and excluded from the non-brain-dominating isoform, then the described switch may lead to differential p53 stability between the brain and other organs by affecting the ZNF500-p53 dimer.

*CAMTA2* regulates cardiac growth (136) and acts in brain development (137). According to our analysis, this gene produces a DBD^+^/DBD^+^ major isoform switch between the brain, as well as the pituitary gland, and many non-brain tissues, along with the cervical spinal cord **(Figure 6E, S9; Table S9)**. While its brain-dominating major isoform has two protein-binding domains (an IQ-motif / EF-hand binding site and an ankyrin repeat-containing domain), its non-brain-dominating major isoform has a catalytic P-loop containing NTP hydrolase domain instead of the IQ-motif **(Table S9)**. Likely, this domain exchange modulates both the binding of partner proteins by CAMTA2 and its catalytic activity which may arise only in non-brain tissues. As CAMTA2 binds calmodulin, which mediates Ca2+ signalling (138), and because the IQ motif is a known calmodulin binder (139), the exchange of the IQ motif for the P-loop containing catalytic domain in the non-brain-dominating major isoform could differentially regulate CAMTA2-dependent Ca2+ signalling within the brain (where only the IQ-motif-containing isoform is expressed) and in other organs (where both isoforms are expressed). Apart from the difference in the coding sequence, the two mRNA isoforms differ by the presence of the 5’-UTR. Only the brain-dominating major isoform has it, and it likely gives additional means of regulation of the stability, translational efficiency or cellular localisation of this mRNA.

Our final example presents a brain-to-non-brain DBD^+^/DBD^+^ major isoform switch produced by *ZBTB45* **(Figure 6F, S10A; Table S9)**, a key regulator of glial differentiation (140). In this switch, the two major mRNA isoforms differ only in their 5’-UTRs **(Table S9)**, due to the usage of alternative transcription start sites. Specifically, the brain-dominating isoform has a longer and completely different 5’-UTR sequence. The two 5’-UTRs could differentially affect the stability, translational efficiency or cellular localisation of the two isoforms. If they are coexpressed in the same cells, then the abundance of ZBTB45 may depend on the relative expression levels of the two isoforms which inverts between the brain and non-brain tissues **(Figure 6F)**. Additionally, TF genes *CEBPG*, *ZC3H8*, *ZNF202* and *ZNF569*, which have not been characterised in the normal brain physiology, also demonstrate brain-to-non-brain switches of DBD^+^ major isoforms that differ only by UTR sequences **(Figure S10B-E; Table S9)**. This fact calls for experimental investigations probing the four TFs in brain cell types.

In total, we predicted and interpreted a set of uncharacterized DBD^+^/DBD^−^ and DBD^+^/DBD^+^ major isoform switches between the brain and other organs. Based on the predicted effects of these switches and the lack of the brain-specific functional knowledge about some of the TF genes that produce them, we suggest that these switches could be experimentally verified and, if confirmed, functionally investigated.

## Discussion

Here, we computationally characterised, for the first time, the alternative isoforms of the full set of known human TF genes in the normal physiology. We found more than 500 major isoform switches between adult human tissues, including 45 pronounced switches between the brain and non-brain tissues, and predicted their functional consequences. We presented detailed interpretations of particular brain-to-non-brain switches, motivating their experimental verification. Moreover, we revealed the scale of DBD^−^ isoform production across adult human tissues, assigned a major part of DBD^−^ isoforms to general functional classes and highlighted broadly and highly expressed DBD^−^ isoforms. To ground our interpretations of isoform domain architectures and to facilitate future studies of TF isoforms, we created a compendium of 92 TF genes (or small groups of closely related TF genes) whose alternative isoforms have been characterised to a varying extent. As TF isoforms have been studied mostly individually over several decades, our collection of references to as many examples as we were able to find in literature will hopefully serve as a starting point for future investigations.

Relative expression levels of major isoforms across normal adult human tissues revealed 536 isoform switches produced by 24% (360/1,476) of expressed TF genes, which suggests that such major isoform switches may represent a widespread, yet understudied, regulatory mechanism. Switch-like isoform expression between tissues in normal physiology have been known in mammals for a long time (2, 141–145). However, isoform switches produced by human TF genes across adult normal tissues have been so far represented only by a handful of examples. Therefore, our collection of cross-tissue major isoform switches with predicted general functional consequences holds a potential to greatly expand the established switch set. As our definition of isoform switches allows considerable coexpression of switching isoforms within tissues, the switches that we predicted may differentially regulate the combined effect of coexpressed alternative proteins.

Interestingly, 48% of switches produced a change outside domains or even outside the coding sequence. Changes in the coding sequence between structured domains may affect intrinsically disordered regions (IDRs) which were previously found to be enriched in alternative exons and regulate protein-protein interactions (PPIs) in a tissue-specific manner (16, 146). Apart from mediating PPIs (147–152), IDRs facilitate DNA binding (152, 153) and RNA binding (154), suggesting that major isoform switches producing non-domain changes in the protein sequence have a potential to rewire a vast network of TF interactions. In the case of differences only in UTRs, a switch may affect the stability, cellular localisation or translational efficiency of the major mRNA isoform (155–157) and, in this way, differentially regulate the production of the corresponding protein. In the light of this established evidence, our observation about the frequency of changes outside domains and outside the coding sequence between switching major isoforms suggests the importance of the verification and further study of these non-domain switches, even taking into account the likely incompleteness of our domain annotation.

Switch clustering by tissue specificity revealed prominent patterns of major isoform exchange between the brain and non-brain tissues. This finding is in line with the previous evidence of a unique alternative splicing repertoire of the brain (158–160) and with the discovery of mammalian brain-to-non-brain isoform switches produced by alternative splicing of microexons (142). Of note, Merkin and colleagues also found that the combination of the heart and muscle tissues has distinct alternative splicing signatures (160), in accord with our observation that the heart and skeletal muscles cluster together when using the complete linkage. Further supporting our results, Tapial and co-authors observed distinct alternative splicing profiles of neural and muscle tissues, including the heart (13). However, to the best of our knowledge, the only experimentally discovered brain-to-non-brain isoform switches produced by TF genes in normal physiology are those by *IRF3* (104), *MEF2A* and *MEF2D* (105), *MEF2C* (106), *TEAD1* (109) and, less clearly, by *HNF4A* (103). Therefore, our collection of 45 unique predicted brain-to-non-brain switches from the complete linkage (CL) cluster 5 and Ward’s D2 linkage (WD2L) cluster 3 is of a particular interest. To motivate future investigations, we concluded our analysis with detailed mechanistic and functional interpretations of particular brain-to-non-brain major isoform switches.

Major isoform switches described in our study suggest experimental verification at the level of mRNA and protein. It could be done by detecting the differential major status of the switching isoforms between two relevant cell lines or primary cell types using real-time PCR with isoform-specific primers and TF-specific quantitative Western blot experiments. Next, single-cell long-read transcriptomics based on the Oxford Nanopore (161–167), or PacBio (168–174), or both of these sequencing technologies (175, 176), as well as single-cell proteomics [reviewed in (177)], would allow determining if switching isoforms are coexpressed in the same cells or produced by different cell subpopulations. This, in turn, would inform switch interpretation and further experimental efforts.

Our ability to analyse DBD^−^ isoforms (**Supplementary Discussion**) and to interpret major isoform switches was limited by several technical and biological factors. First one is an incomplete representation of transcription regulation domains in the InterPro database, along with the lack of functional annotation for some domains both in the database and in literature. Also, our reductionist approach of assigning one particular function to each domain may not always be adequate, as some domains are known to play multiple functional roles (178). Additionally, our assumption that DBD^−^ isoforms do not facilitate the same function as their DBD^+^ counterparts cannot always hold, as it has been proven wrong for particular isoforms (90, 179). Of note, we do not assume that every DBD^−^ or DBD^+^ isoform has a particular function, but instead predict a likely functional class of each isoform, in case it does have a function *in vivo*. On the other hand, we may have underestimated the number of TF isoforms that cannot bind DNA, as we assumed that all DBD^+^ isoforms have the potential to do so. On the contrary, C2H2 ZF isoforms may lose the DNA-binding ability even when still containing some of their zinc fingers (180). Also, TFs that dimerize to bind DNA, such as the members of the bHLH and bZIP families, would lose DNA binding upon losing the dimerization domain.

More generally, by the design of the study, we did not take into account certain mechanisms of proteome diversification, such as alternative translation of the same mRNA isoform (181–184) and post-translational modifications (PTMs) affecting protein isoform properties (185). Finally, due to the key role of protein domains in our analysis, we were not able to interpret differences in IDRs and excluded non-coding RNA isoforms produced by TF genes, although some non-coding TF isoforms have been proven functionally important (186–188). Overall, due to the incomplete domain repertoire, potential alternative translation and PTMs, as well as the exclusion of IDR differences and non-coding isoforms, we may have underestimated the potential functional significance of alternative TF isoforms and their switches.

In spite of the described limitations, our results demonstrate the importance of studying TF genes at the isoform level and suggest that major isoform switches by human TF genes represent a widespread and as yet understudied level of transcription regulation.

## Data availability

The data underlying this article are available in the article, in its online supplementary material and in Zenodo, at https://zenodo.org/records/14181731 (Supplementary Data File, DOI: 10.5281/zenodo.14181730).

## Acknowledgements

We would like to thank our colleagues from the Computational Regulatory Genomics Group and the Bioinformatics and Computational Biology Laboratory for fruitful discussions and feedback.

## Contributions

N.M.L., S.S. and B.L.: Project conceptualization. S.S. and K.P.: Investigation and data analysis. S.S.: Writing. N.M.L. and B.L.: Supervision, review and editing, funding acquisition.

