## Supplementary Text and Figures for "Cross-tissue isoform switches by human transcription factors suggest a widespread regulatory mechanism"

### Supplementary Discussion

We began our analysis by considering isoforms that lack a DNA-binding domain (DBD$^{-}$ isoforms) and hence cannot, by definition, serve as TFs. In their pioneering study, Resch and colleagues reported that several DBD types, including C4 zinc fingers, SANT and homeobox domains, exhibit particularly high levels of alternative splicing in humans (1). In line with this evidence, Taneri and co-authors demonstrated that DBDs comprise 75% of all alternatively spliced murine domains with known functions (2), while later on Talavera and colleagues showed that alternative splicing frequently changes DBDs in human and mouse (3). Much more recently, Soto and co-authors found that effector domains more frequently differ in alternative isoforms of human TF genes than DBDs; however, they also reported that 42 TFs had a DBD affected in the majority of proteoforms (4). Here, we expand these analyses by revealing the scale of the DBD$^{-}$ isoform production by human TF genes, based on the high-quality subset of coding transcripts from a recent human genome annotation. We found that almost a third of human TF genes have at least one DBD$^{-}$ isoform in the genome annotation and that such isoforms comprise 17% of all isoforms annotated for human TF genes. Moreover, we have shown that 69% of all annotated DBD$^{-}$ isoforms are expressed in at least one adult human tissue, while 21% of all annotated DBD$^{-}$ isoforms become the highest-expressed (major), in comparison to other isoforms of the same TF gene, in at least one tissue. Together, these findings suggest a prominent role of DBD$^{-}$ isoforms in human adult tissues, depending on the level of their translation which needs to be investigated.

Our tissue-specificity analysis of DBD$^{-}$ isoforms revealed more than 30 broadly expressed DBD$^{-}$ isoforms that are produced at a high level in at least some adult human tissues. Importantly, four such isoforms that we selected for an in-depth description are coexpressed, at the level of the whole tissues, with one or more DBD+ isoforms produced by the same TF gene across the whole panel of studied tissues. This fact suggests a combined action of the broadly expressed DBD+ and DBD$^{-}$ isoforms, if they are coexpressed in individual cells. Indeed, Tapial and co-authors demonstrated that >2,600 human genes coexpress at least two isoforms across the majority of tissues and cell types that they studied and that the coexpression of alternative mRNA isoforms occurs predominantly within individual cells (5). In addition, Weatheritt and colleagues have shown that the majority of alternative isoforms are translated (6). Crucially, Tapial and co-authors also found that the set of broadly expressed genes producing at least two mRNA isoforms per tissue is enriched in genes involved in transcription regulation and chromatin modification (5). Therefore, together with these previous findings, our observations suggest a functional interplay between DBD$^{-}$ and DBD+ isoforms coexpressed in individual cells at the protein level and broadly expressed across adult human tissues.

### Supplementary Figures

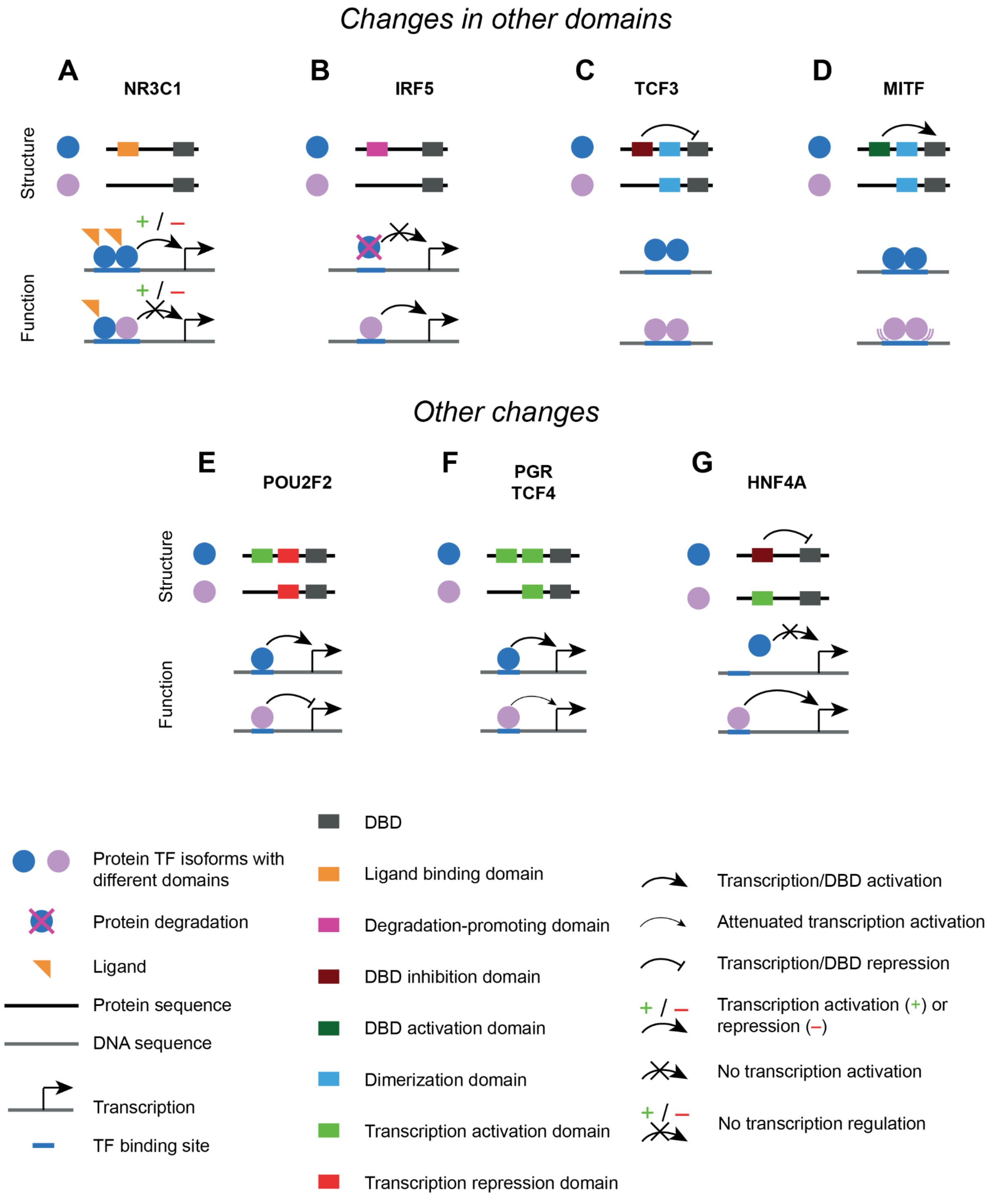


**Figure S1. Differences in domains between alternative isoforms affect the function of a TF gene. (A-D)** Differences in domains other than DBDs or transcription regulation domains. **(E-G)** More complex differences in transcription regulation domains, than those presented in **Figure 1**. See **Table S1** for details.

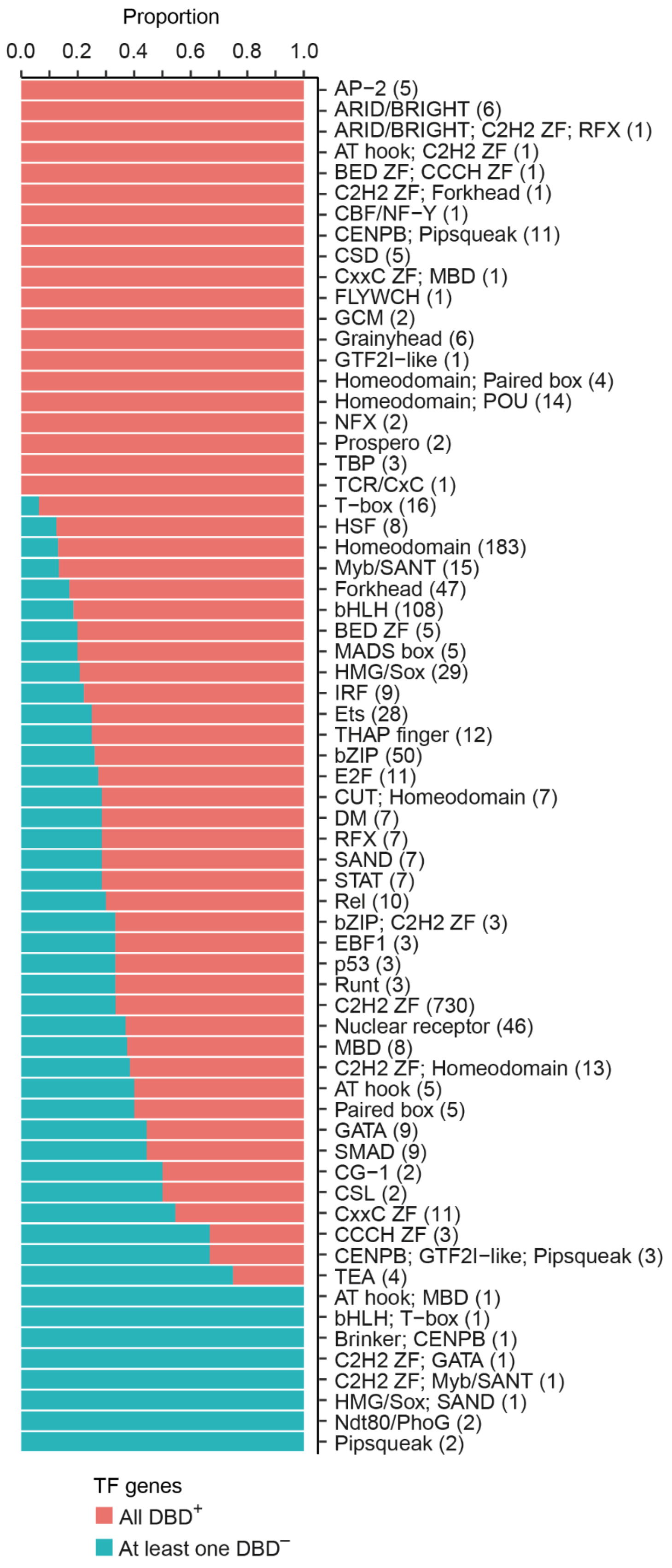


**Figure S2. Proportions of TF genes that have only DBD$^+$ isoforms (red) or at least one DBD$^-$ isoform (blue), per structural family, based on the whole-genome annotation.**

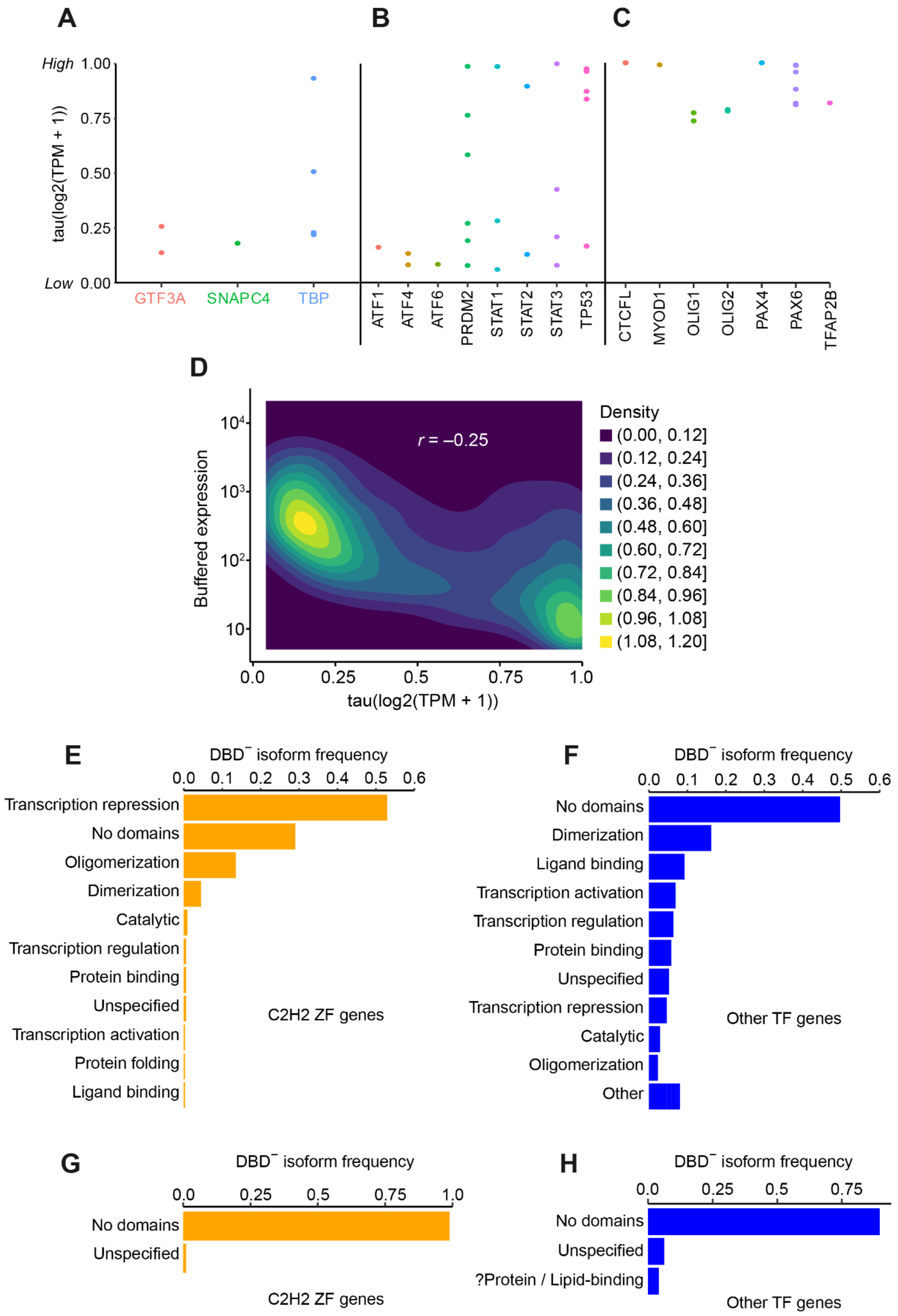

A
B
C
High 1.00
0.75
0.50
0.25
Low 0.00
tau(log2(TPM + 1))
GTF3A
SNAPC4
TBP
ATF1
ATF4
ATF6
PRDM2
STAT1
STAT2
STAT3
TP53
CTCFL
MYOD1
OLIG1
OLIG2
PAX4
PAX6
TFAP2B
D
r = –0.25
Buffered expression
10^4
10^3
10^2
10
0.0
0.25
0.5
0.75
1.0
tau(log2(TPM + 1))
Density
(0.00, 0.12]
(0.12, 0.24]
(0.24, 0.36]
(0.36, 0.48]
(0.48, 0.60]
(0.60, 0.72]
(0.72, 0.84]
(0.84, 0.96]
(0.96, 1.08]
(1.08, 1.20]
E
DBD⁻ isoform frequency
0.0 0.1 0.2 0.3 0.4 0.5 0.6
Transcription repression
No domains
Oligomerization
Dimerization
Catalytic
Transcription regulation
Protein binding
Unspecified
Transcription activation
Protein folding
Ligand binding
C2H2 ZF genes
F
DBD⁻ isoform frequency
0.0 0.1 0.2 0.3 0.4 0.5 0.6
No domains
Dimerization
Ligand binding
Transcription activation
Transcription regulation
Protein binding
Unspecified
Transcription repression
Catalytic
Oligomerization
Other
Other TF genes
G
DBD⁻ isoform frequency
0.0 0.25 0.5 0.75 1.0
No domains
Unspecified
C2H2 ZF genes
H
DBD⁻ isoform frequency
0.0 0.25 0.5 0.75
No domains
Unspecified
?Protein / Lipid-binding
Other TF genes

**Figure S3. Tissue specificity (measured with the tau index) and predicted functional classes of the expressed isoforms of human TF genes.** Tissue specificity of the isoforms of general transcription factor (GTF) subunits **(A)**, broadly expressed TFs **(B)** and tissue-specific TFs **(C)**, calculated to test the efficiency of the tau index as a measure of tissue specificity. Panels **A** and **B** demonstrate that GTF subunits and broadly expressed TFs each have at least one tissue-non-specific isoform, as expected. Panel **C** shows that all expressed isoforms of tissue-specific TFs are highly tissue-specific, as expected. TFs in panels **B** and **C** were selected according to the gene-level single-cell transcriptional profiles from the Human Protein Atlas (proteinatlas.org). **(D)** Density of expressed isoforms ($N$ = 3,401) of TF genes as a function of the buffered isoform expression level and isoform tissue specificity. **(E-F)** Proportions of $DBD^-$ isoforms expressed by C2H2 ZF genes ($N$ = 310 $DBD^-$ isoforms) **(E)** or other TF genes ($N$ = 173 $DBD^-$ isoforms) **(F)** per functional class of a non-DBD contained in a $DBD^-$ isoform. As a $DBD^-$ isoform may contain more than one functional class of non-DBDs, the proportions may not sum to 1. **(G, H)** Proportions of expressed $DBD^-$ isoforms assigned to the "Unclear" functional class **(Figure 2F-G)** and produced by C2H2 ZF genes **(G)** or other TF genes **(H)**. The isoforms are stratified by the presence and functional class of their non-DBDs. The "Unspecified" group of non-DBDs contains domains to which we were not able to assign a function.

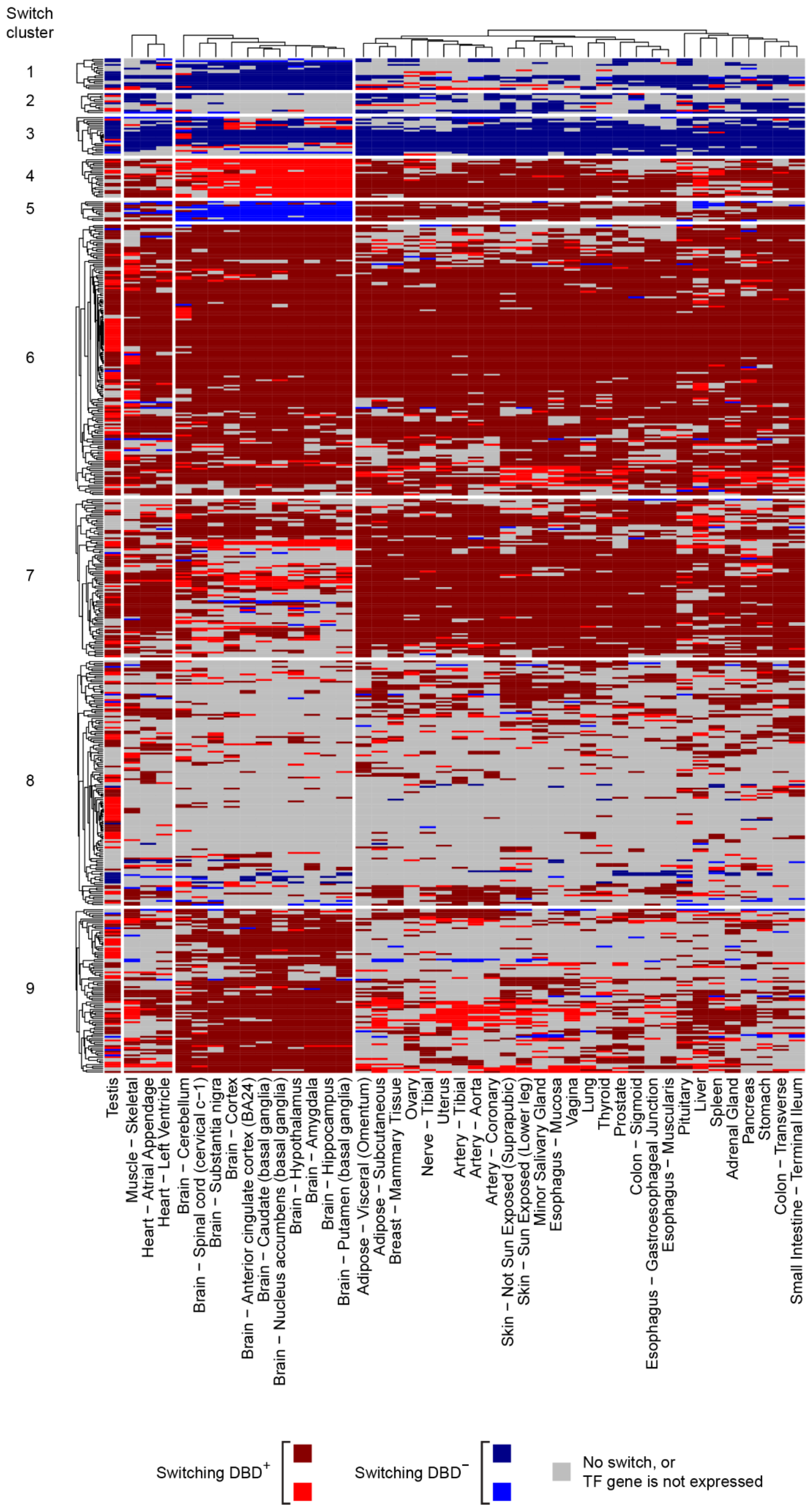


**Figure S4. Major isoform switches (rows) across adult human tissues (columns).** See **Figure 5A** for the description of the colour scheme. The heatmap is clustered using the complete linkage, according to switch expression patterns.

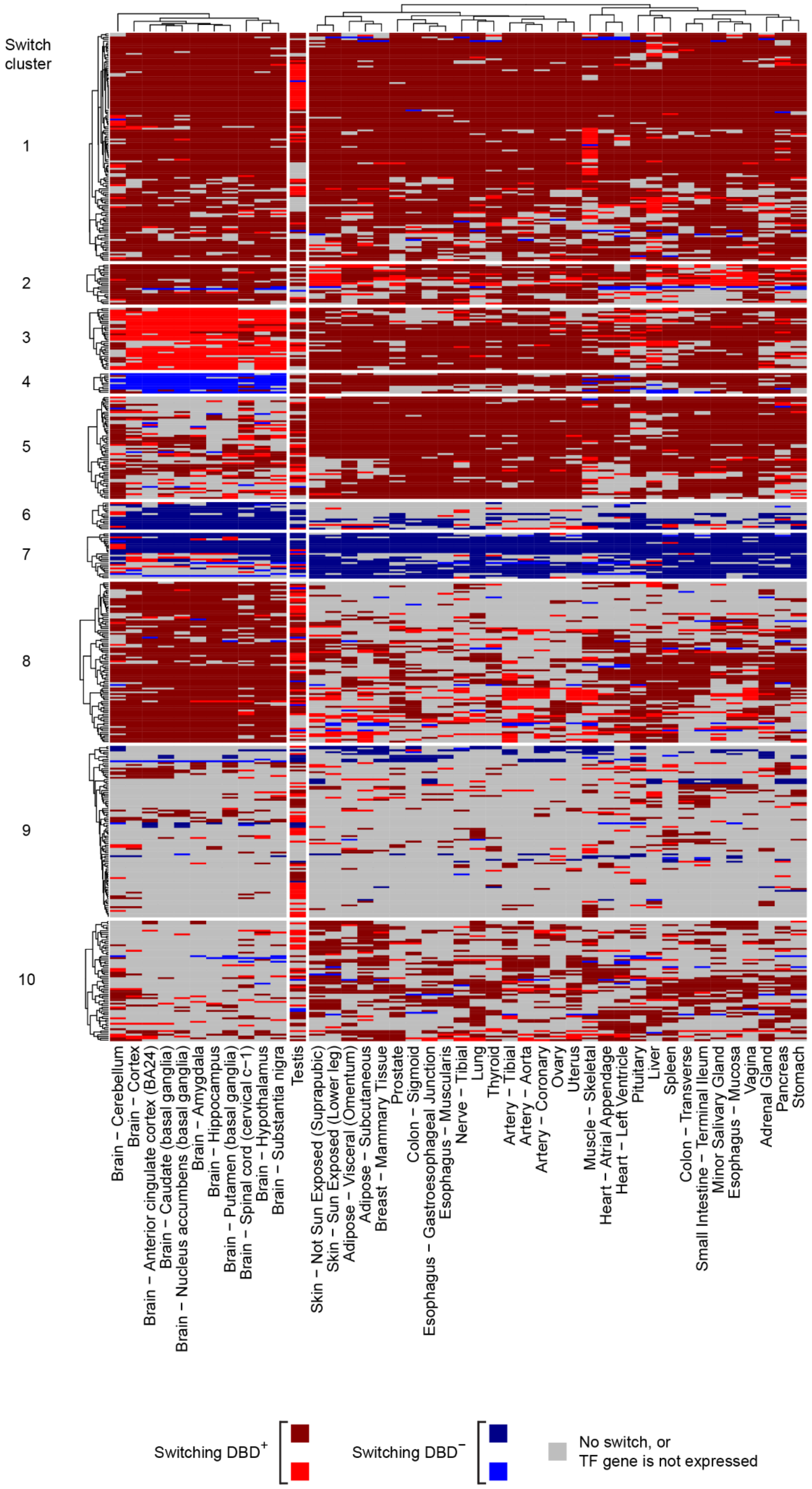


**Figure S5. Major isoform switches (rows) across adult human tissues (columns).** See **Figure 5A** for the description of the colour scheme. The heatmap is clustered using Ward's D2 linkage, according to switch expression patterns.

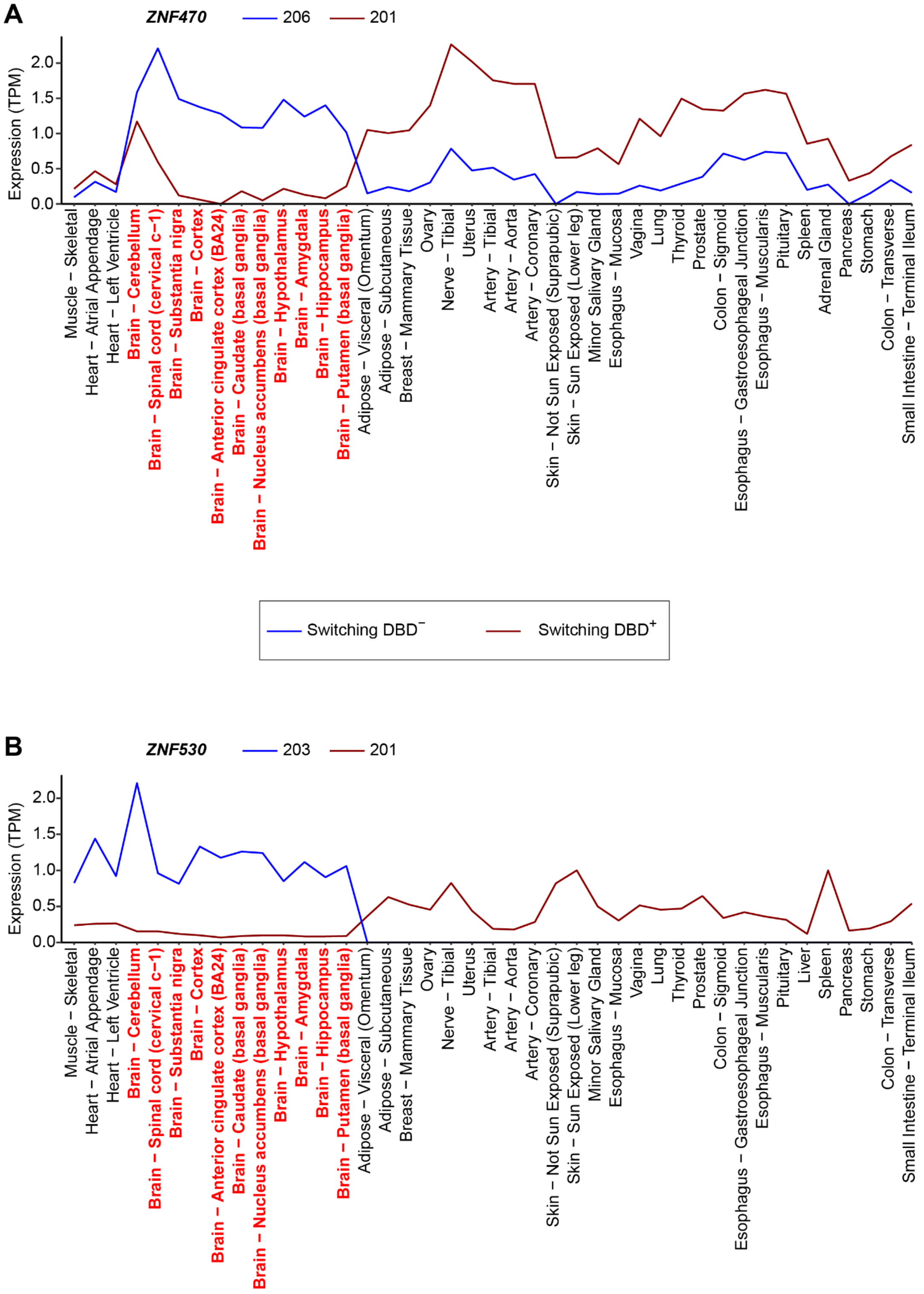


**Figure S6. DBD$^+$/DBD$^−$ major isoform switches produced by TF genes *ZNF470* (A) and *ZNF530* (B) between the brain and non-brain tissues.** The names of the brain tissues are highlighted in red. See **Figure 5A** for the description of the colour scheme for switching isoforms. Isoforms are identified by their numbers in the Ensembl v99 human genome annotation. For instance, isoform 206 in panel **A** is *ZNF470*-206 (ENST00000601902). (Continued below.)

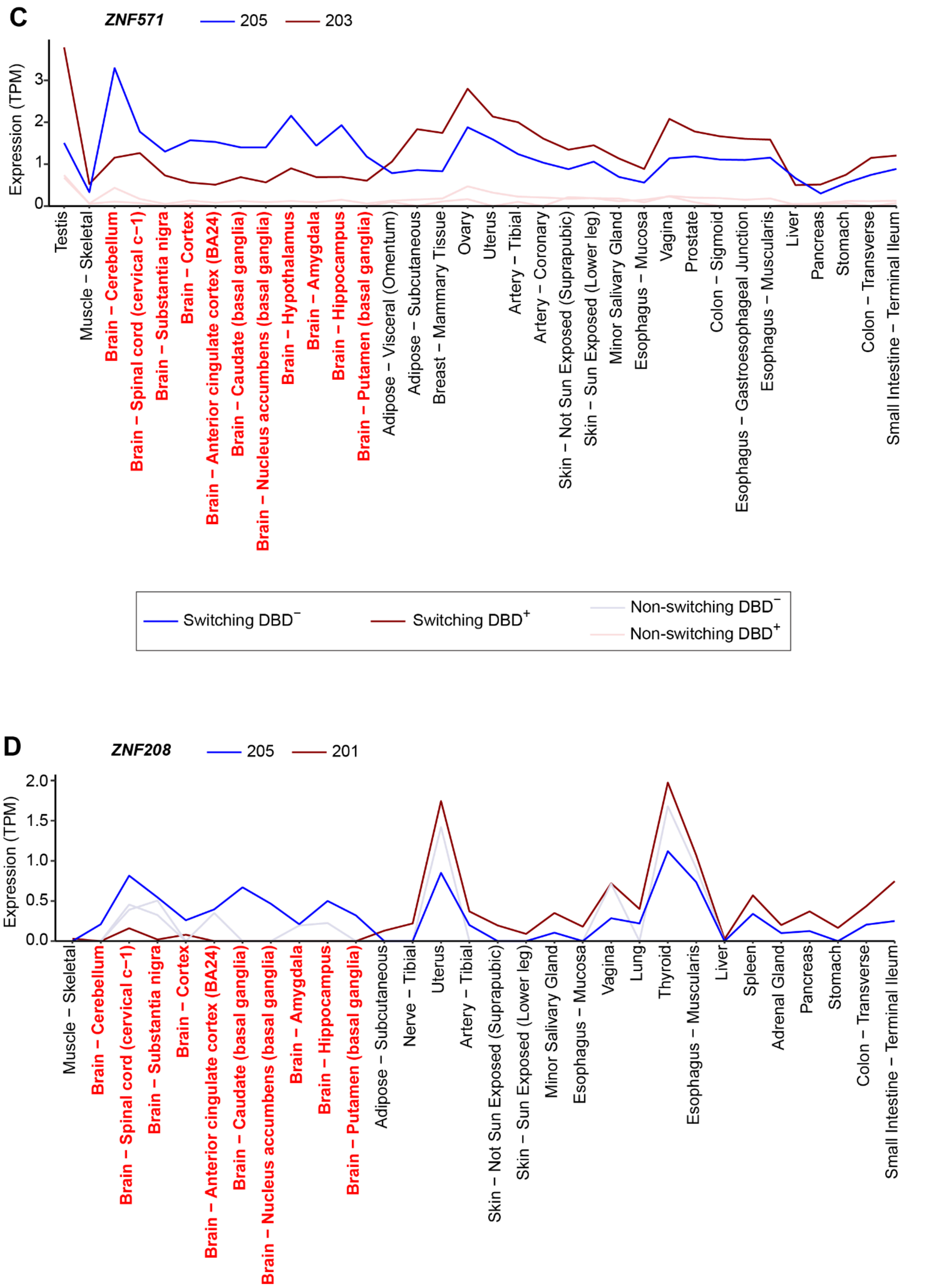


**Figure S6 (continued). DBD$^{+}$/DBD$^{-}$ major isoform switches produced by TF genes *ZNF571* (C) and *ZNF208* (D) between the brain and non-brain tissues.**

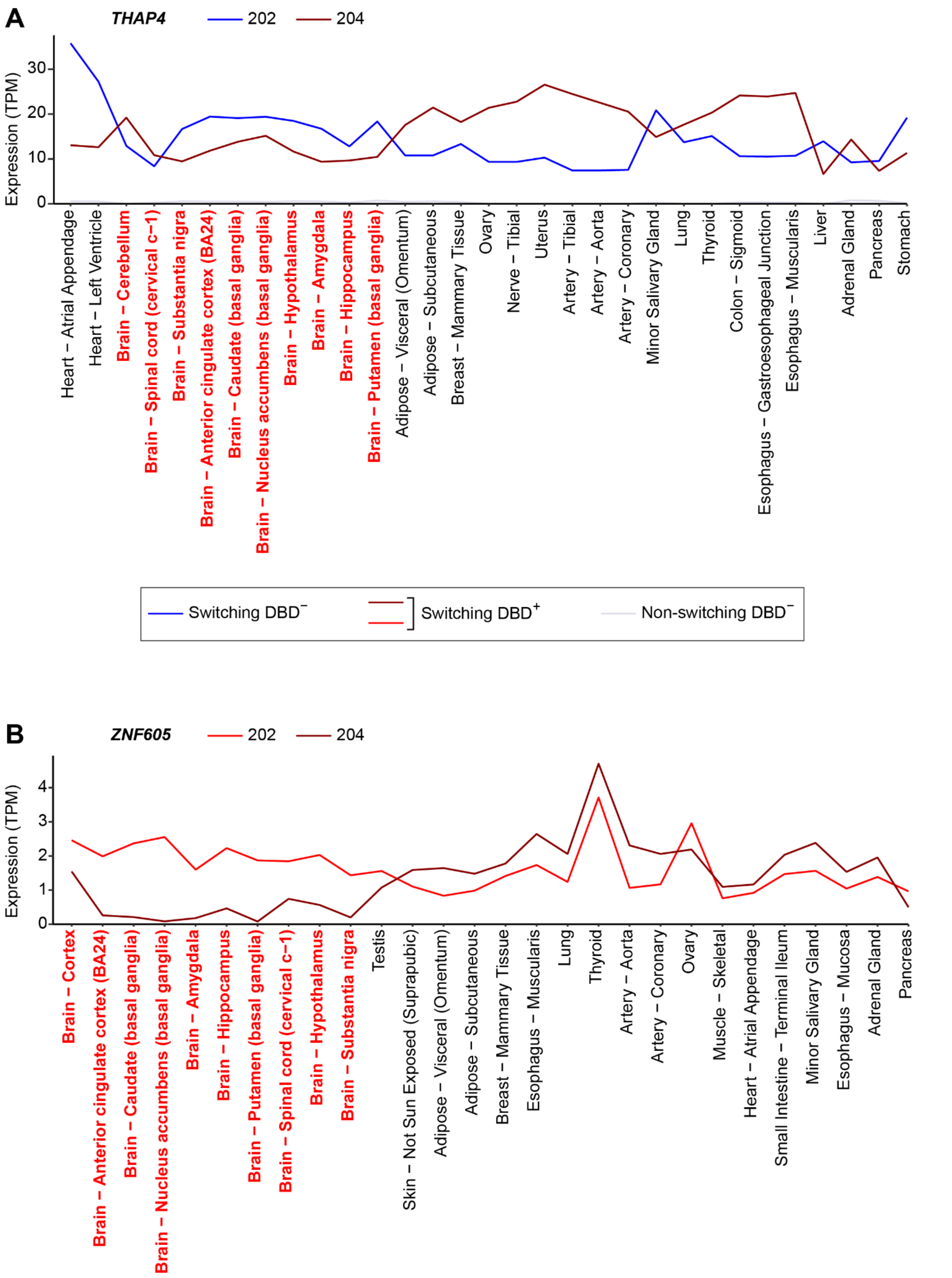


**Figure S7. Major isoform switches produced by TF genes *THAP4* (A) and *ZNF605* (B) between the brain and non-brain tissues.** The names of the brain tissues are highlighted in red. See **Figure 5A** for the description of the colour scheme for switching isoforms. Isoforms are identified by their numbers in the Ensembl v99 human genome annotation. For instance, isoform 202 in panel **A** is *THAP4*-202 (ENST00000402136).

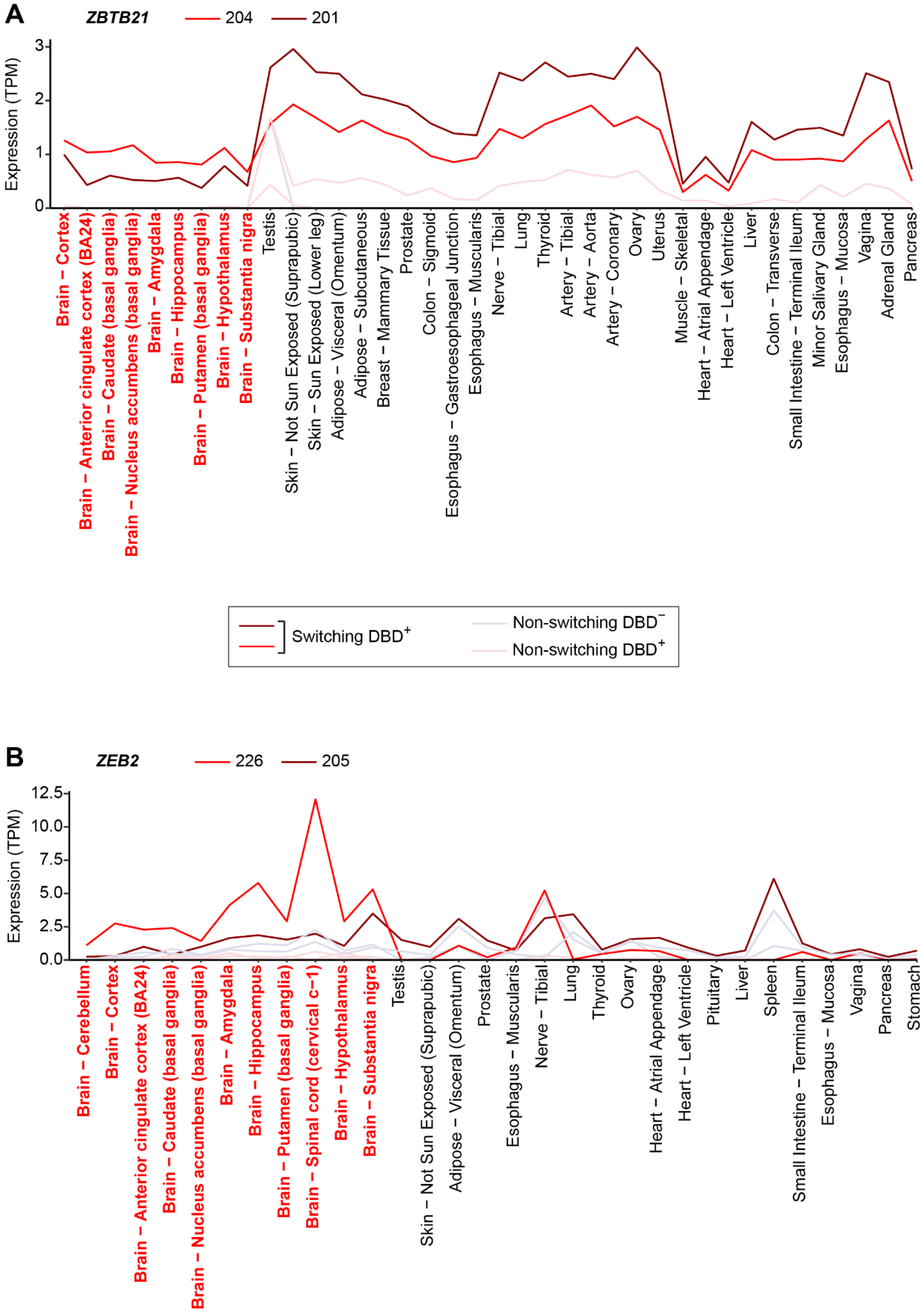


**Figure S8. $DBD^+$/$DBD^+$ major isoform switches produced by TF genes *ZBTB21* (A) and *ZEB2* (B) between the brain and non-brain tissues.** The names of the brain tissues are highlighted in red. See **Figure 5A** for the description of the colour scheme for switching isoforms. Isoforms are identified by their numbers in the Ensembl v99 human genome annotation. For instance, isoform 204 in panel **A** is *ZBTB21*-204 (ENST00000398505). (Continued below.)

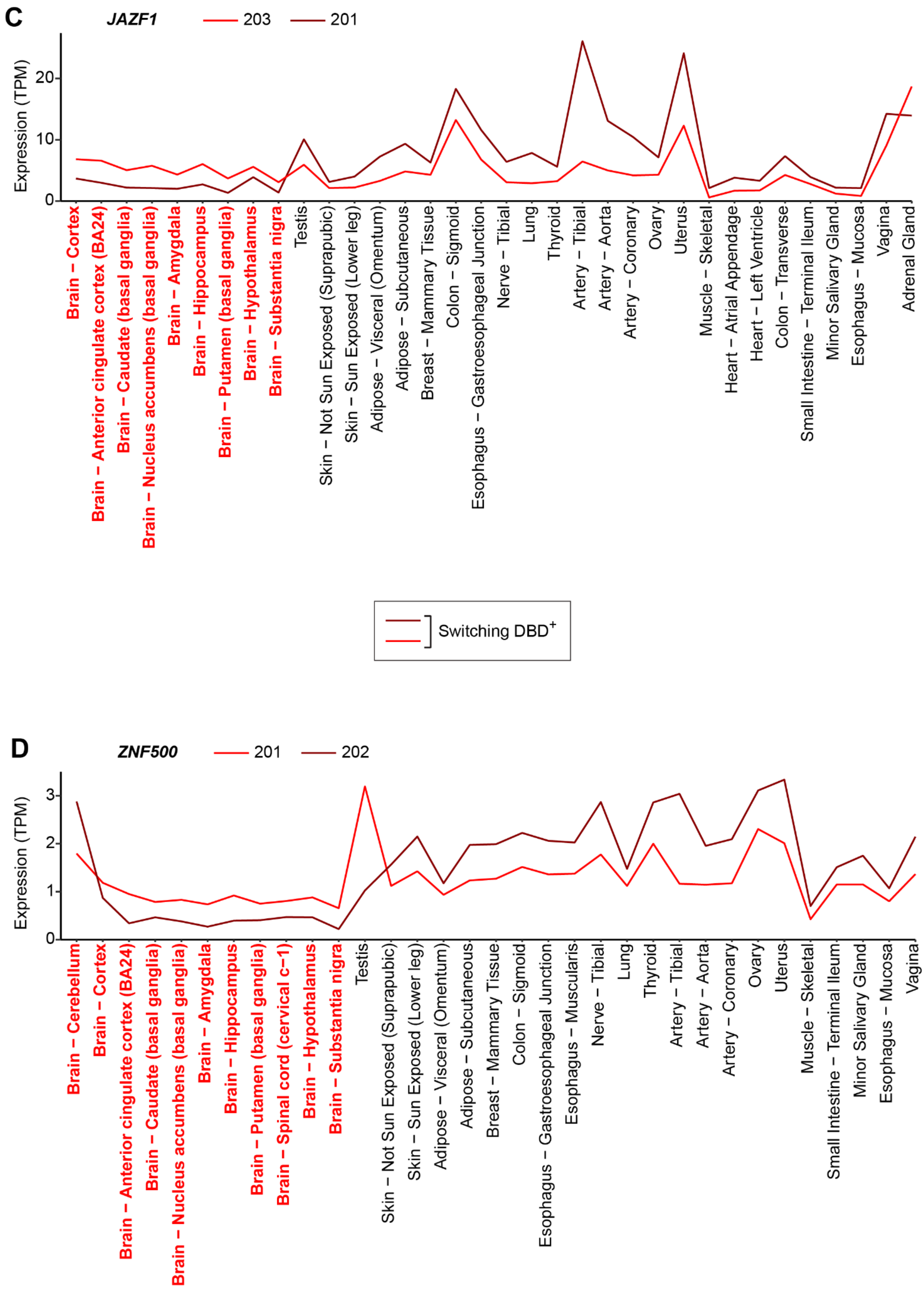


**Figure S8 (continued). DBD$^+$/DBD$^+$ major isoform switches produced by TF genes *JAZF1* (C) and *ZNF500* (D) between the brain and non-brain tissues.**

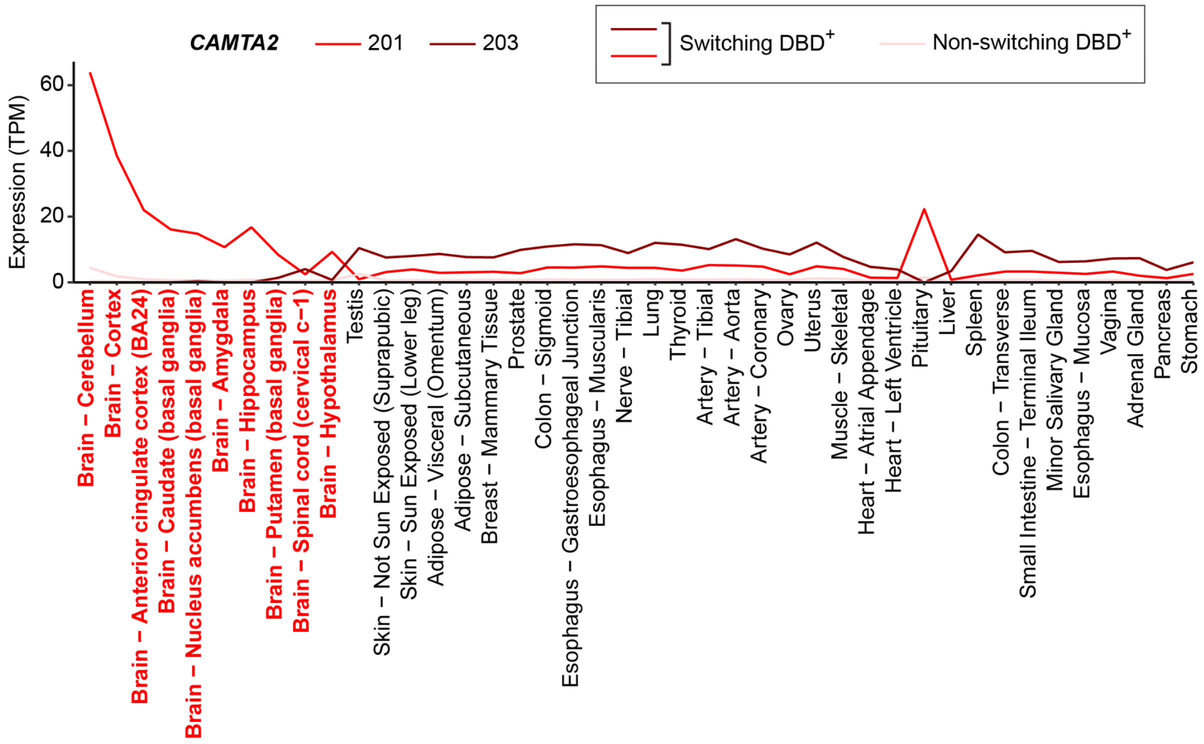


**Figure S9. A DBD+/DBD+ major isoform switch produced by a TF gene *CAMTA2* between the brain and non-brain tissues.** The names of the brain tissues are highlighted in red. See **Figure 5A** for the description of the colour scheme for switching isoforms. Isoforms are identified by their numbers in the Ensembl v99 human genome annotation. For instance, isoform 201 is *CAMTA2*-201 (ENST00000348066).

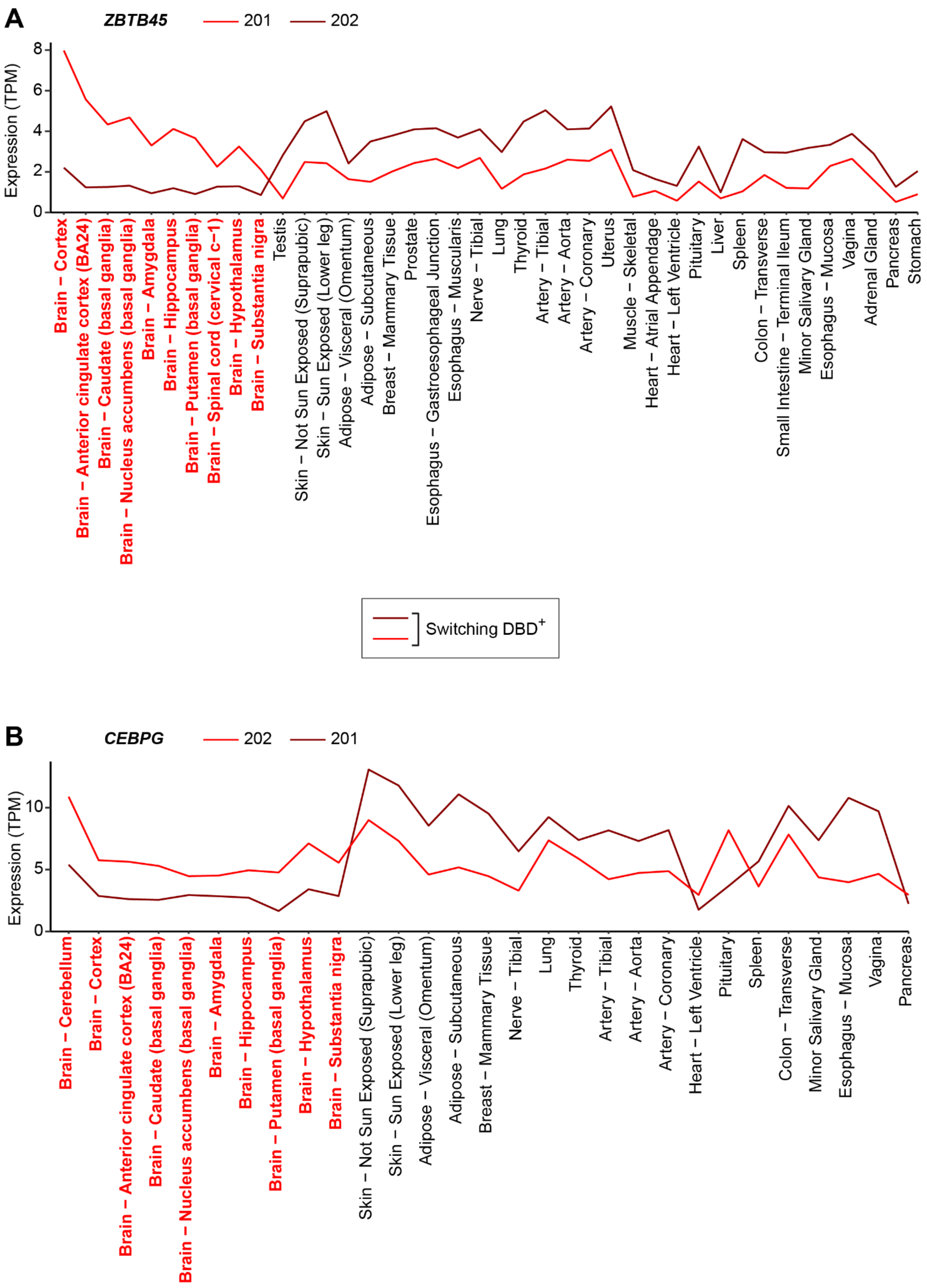


**Figure S10. DBD$^+$/DBD$^+$ major isoform switches produced by TF genes *ZBTB45* (A) and *CEBPG* (B) between the brain and non-brain tissues.** The names of the brain tissues are highlighted in red. See **Figure 5A** for the description of the colour scheme for switching isoforms. Isoforms are identified by their numbers in the Ensembl v99 human genome annotation. For instance, isoform 201 in panel **A** is *ZBTB45*-201 (ENST00000354590). (Continued below.)

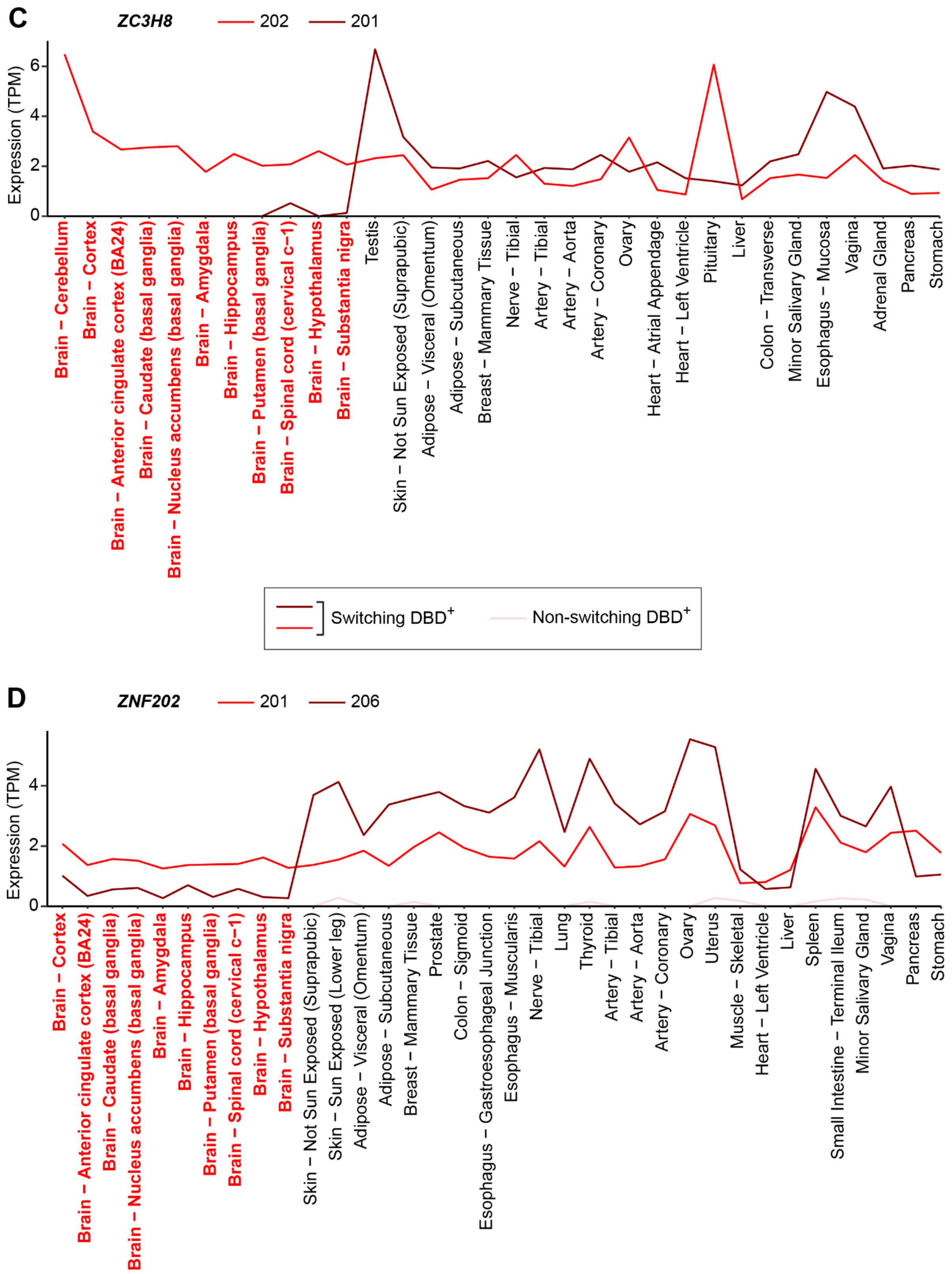


**Figure S10 (continued). DBD$^+$/DBD$^+$ major isoform switches produced by TF genes *ZC3H8* (C) and *ZNF202* (D) between the brain and non-brain tissues. (Continued below.)**

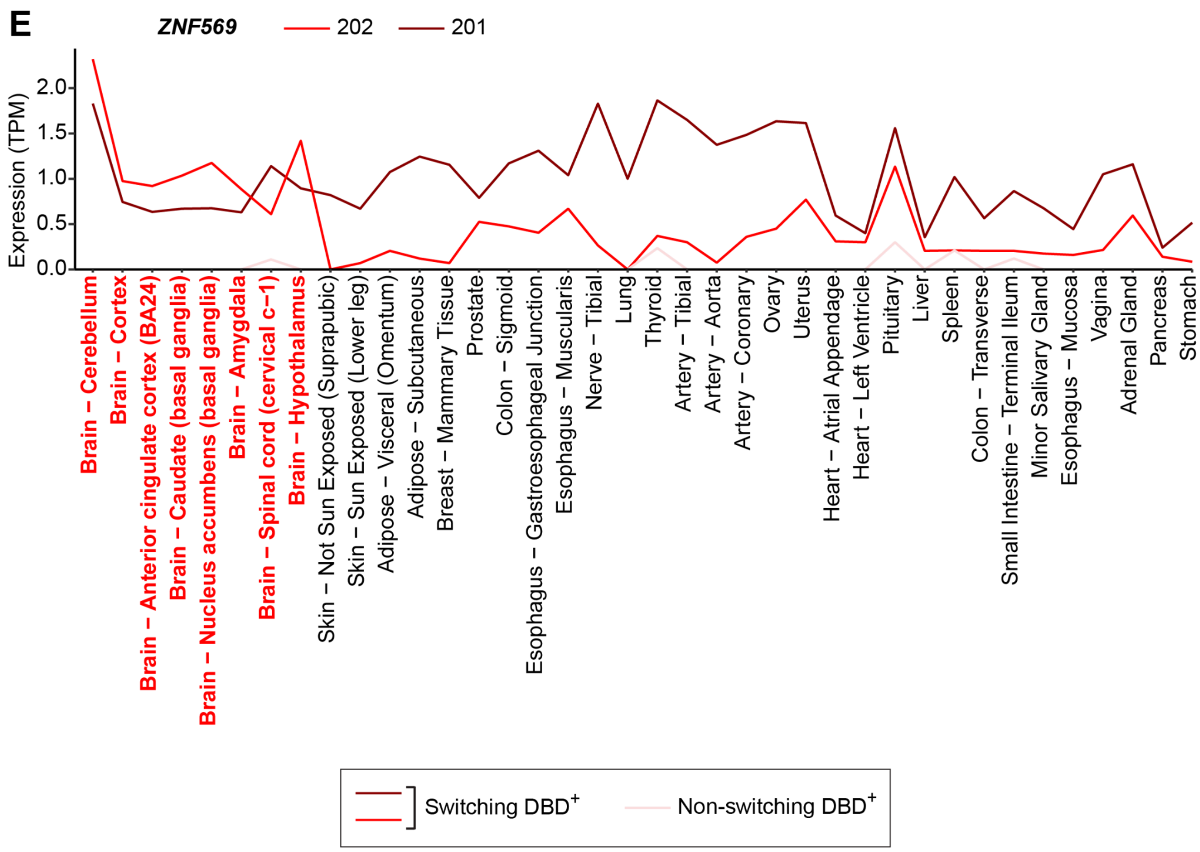


**Figure S10 (continued). A DBD+/DBD+ major isoform switch produced by a TF gene *ZNF569* (E) between the brain and non-brain tissues.**
